# Urolithin A improves Alzheimer’s disease cognition and restores mitophagy and lysosomal functions

**DOI:** 10.1101/2024.01.30.577986

**Authors:** Yujun Hou, Xixia Chu, Jae-Hyeon Park, Qing Zhu, Mansoor Hussain, Zhiquan Li, Helena Borland Madsen, Beimeng Yang, Yong Wei, Yue Wang, Evandro F. Fang, Deborah L. Croteau, Vilhelm A. Bohr

## Abstract

**Background:** Compromised autophagy, including impaired mitophagy and lysosomal function, is thought to play a pivotal role in Alzheimer’s disease (AD). Urolithin A (UA) is a gut microbial metabolite of ellagic acid that stimulates mitophagy. The effects of early and/or long-term treatment, as well as more detailed mechanisms of action, are not known.

**Methods:** We addressed these questions in three mouse models of AD, and behavioral, electrophysiological and biochemistry assays were performed.

**Results:** Long-term UA treatment significantly improved learning, memory and olfactory function in different AD transgenic mice. UA also reduced Aβ and Tau pathologies, and improved long-term potentiation. We found that UA activated autophagy/mitophagy via increasing lysosomal functions. At the cellular level, UA improved lysosomal function and normalized lysosomal cathepsins, especially targeting cathepsin Z, to restore lysosomal function in AD, indicating the important role of cathepsins in UA-induced therapeutic effects of AD.

**Conclusions:** Collectively, our study highlights the importance of lysosomal dysfunction in AD etiology, and points to the high translational potential of UA.

## 1. INTRODUCTION

Alzheimer’s disease (AD) is a neurodegeneration that affects over 10% of adults age 65 and older and its prevalence is estimated to triple worldwide by 2050, a huge burden in our aging society [1, 2]. The clinical manifestations include cognitive impairments and abnormal behaviors. The main classical pathological features of AD are amyloid plaques formed by amyloid beta (Aβ) and neurofibrillary tangles formed by phosphorylated tau [3]. Aging is a major risk factor for neurodegenerative diseases, and many aging hallmarks play important roles in the pathogenesis and development of AD, including compromised autophagy [4], DNA damage, neuroinflammation, cellular senescence, and mitochondrial dysfunction [5–7]. Autophagy is markedly impaired in AD and autolysosome acidification in AD mouse models induces autophagic build-up of Aβ in neurons, yielding senile plaques, thus compromised lysosomal function is thought to be a driver of AD [4].

Mitophagy is a degradation process that clears damaged or superfluous mitochondria. The timely removal of damaged mitochondria plays an essential role in maintaining normal physiological function and is vital for the survival and health of neurons [8, 9]. Studies by us and others have shown that mitophagy levels in the brains of AD patients are compromised and that mitophagy in transgenic mice, nematode models of AD, and iPSC-derived neurons of AD patients are impaired [10, 11]. Mitophagy induction by genetic strategies or small molecular compounds can improve the cognitive function of AD mice and reduce the level of Aβ plaques and phosphorylated Tau (pTau) in brains [11, 12]. Many natural mitophagy inducers could reduce AD symptoms, including Urolithin A (UA), nicotinamide riboside (NR), Kaempferol, and Rhapontigenin etc. [11, 13–15].

UA is a natural compound produced by gut bacteria that ingests ellagitannins and ellagic acid, and is a complex polyphenol found in foods such as pomegranates, berries, and nuts [16, 17]. UA was discovered 40 years ago [18] and is considered to be the most conserved and studied urolithin across species [16], but only recently has its impact on aging and diseases been explored. UA prolongs the lifespan of *C. elegans* and safeguards against physiological decline, as illustrated by improved muscle function in young animals and the prevention of age-related muscle decline in old mice [19]. UA also enhances cellular health by increasing mitophagy, mitochondrial function and reducing detrimental inflammation [11, 19, 20]. We recently reported that short-term (2 months) UA treatment induced mitophagy in AD mouse brain and in AD nematodes, and that it improved learning and memory in APP/PS1 AD and 3xTgAD mice [11]. Our results were later validated [21]. Mitophagy genes including PTEN-induced kinase 1 (*PINK1*), pleiotropic drug resistance 1 (*PDR1*) and DAF-16/FOXO-controlled germline-tumor affecting-1 (*DCT1*) play important roles in ameliorating memory impairments and prolong the lifespan of *C. elegans* [11, 22]. In accord with this, UA effectively increased the levels of PINK1, PDR1 and DCT1, highlighting the potential application of UA in AD therapy to improve mitochondrial function and health status [20]. UA treatment had anti-neuroinflammatory effects in activated microglia, supporting the potential neuroprotective role of UA in AD brains [23]. Other studies have confirmed the anti-inflammatory properties of UA *in vivo* and *in vitro* [17, 21].

Other mechanisms of action have been proposed for UA, including the activation of the Ahr/Nrf2 pathway and its downstream antioxidative stress response, the removal Aβ from neurons, the inhibition of DYRK1A activity, [24]and the inhibition of regulators of cancer cell proliferation [15, 24, 25]. The contribution of these mechanisms to the positive impact of UA on aging needs further study. UA has so far only been investigated in humans for its benefits on mitochondrial and muscle health [17, 26]. A phase I clinical study confirmed that UA was safe in healthy, sedentary older adults and activation of mitochondrial biomarkers in muscle and plasma was observed, consistent with previous work shown in cells and *in vivo* in model organisms [26]. UA is well-tolerated and play a therapeutic role in the brain as it crosses the blood-brain barrier [27, 28]. Collectively, growing evidence supports that UA effectively targets AD-related neurodegeneration and holds potential as an intervention for this disease.

Given the low success rate of anti-AD drug development, approaches targeting basic aspects of AD pathologies, such as defective mitophagy, hold therapeutic potential. Aging is the greatest risk factor for the development of AD and, importantly, UA is proposed to target multiple aging and AD deficiencies. Previously, we reported on benefits of UA treatment of AD mice for a brief period (2 months) [11], however most AD patients are likely to receive treatment and care for an extended period of time. Thus, here we have used multiple AD mouse models, APP/PS1, 3xTgAD (AD) and 3xTgAD/Polβ^+/-^ (ADP) mice (DNA repair deficient AD mice made by us), and each were treated with UA for 5 months to study the effects of UA treatment. This study strengthens support for the potential application of UA against AD.

## 2 Methods

### 2.1 Mice

All animal experiments were performed and approved by the National Institute on Aging (NIA) Animal Care and Use Committee, and met all relevant ethical regulations (study protocol number 361-ODS-2023). All animals were maintained at the NIA under standard conditions and fed standard animal feed. Male APP/PS1 mice and their WT littermates were used for the experiments. The APP/PS1 mouse strain (stock no. 004462; The Jackson Laboratory) was obtained from Dr. Mark Mattson’s laboratory.

APP/PS1 mice were treated with UA (200 mg/kg/day) by gavage starting at 2 and ending at 7 months, when behavioral and molecular endpoints were assessed. 3xTgAD and 3xTgAD/Polβ^+/-^ mouse strains were generated as described previously [29]. 12-month-old 3xTgAD mice and 3xTgAD/Polβ^+/-^ mice were treated with UA (200 mg/kg/day) by oral gavage for 5 months, ending at 17 months, followed by evaluation of behavioral and molecular endpoints. Both male and female 3xTgAD and 3xTgAD/Polβ^+/-^ mice were used in the analyses. Statistical methods were not used to predetermine sample sizes, but our sample sizes are similar to those reported in previous studies.

### 2.2 Cell culture

HMC3 human microglia cells were purchased from ATCC. HMC3 were cultured in Eagle’s Minimum Essential Medium (Gibco) with 10% fetal bovine serum in a humidified incubator with 5% CO_2_ at 37°C. 293 APP Swedish cells, were maintained in high-glucose DMEM (Gibco) supplemented with 10% fetal bovine serum (FBS, Gibco, #30084) and cultured in a humidified atmosphere with 5% CO_2_ at 37 °C. When HMC3 cells reached 60% confluence, cells were transfected with (20 nM) siRNAs (KeyGEN BioTECH), 4 h later were cocultured with 10 μM Aβ_42_, 30 μM UA, 3 μM CTSZ inhibitor (MCE, #HY-146985) or together for 48 h, diluting the drug stocks in Eagle’s Minimum Essential Medium with 2% fetal bovine serum. When 293-APP_Swe_ cells reached 60% confluence, cells were transfected with (20 nM) siRNAs, 4 h later were cocultured with 30 μM UA, 3 μM CTSZ inhibitor or both together, diluting the stocks in high-glucose DMEM with 2% fetal bovine serum.

### 2.3 Morris Water Maze test

The MWM test was performed as described previously [11]. The device is a circular pool (140 cm diameter) that is filled with water and kept at 22°C. The pool was painted with non-toxic white paint. A transparent platform (12 cm in diameter) was placed 1 cm below the water surface at a fixed position. Mice were trained for 7 days, with 4 trials per day. Each trial lasted 60 s or until the mouse found the platform. If the mouse did not find the platform within the specified time, the experimenter directed the mouse to the platform. After each trial, the mouse was placed on the platform for 30 s. On the 7th day, after the training phase, the platform was removed for a probe trial which lasts 60 s. All parameters were recorded by a video tracking system (ANYmaze, version 4.99; Stoelting).

### 2.4 Y maze

The Y maze spontaneous alternation performance (SAP) test measures the ability to recognize previously explored environments [30]. The maze consisted of three arms (8 × 30 × 15 cm), with an angle of 120 degrees between them. The number of entries and alterations were recorded by the ANY-maze video tracking system. Mice were introduced to the center of the Y maze and left to explore the maze for 10 min. Between trials the arms were cleaned with 70% ethanol solution. SAP is the subsequent entry into a novel arm over the course of 3 entries; the percentage SAP is calculated by the number of actual alternations/(total arm entries − 2) × 100.

### 2.5 Object recognition test

The object recognition tests were performed as described previously [30]. The device was a Plexiglas box with a size of 25 × 25 × 25 cm^3^. During the training phase, the mice could explore two identical objects for 10 min; During the test phase, 1 h after the training, each mouse was returned to the box, which had been modified to contain a familiar object and a novel object. The boxes and objects were cleaned before each test to eliminate scent clues. The automatic video tracking system (ANY-maze) was used to monitor exploration behavior. Exploration time was calculated as the length of time each mouse sniffed or pointed its nose or paws at the object. The ‘recognition index’ refers to the time spent exploring the novel object relative to the time spent exploring both objects.

### 2.6 Olfactory test

The procedure for the buried food test has been described previously [31]. Mice were fasted for 16 h and then placed in a cage containing bedding that was 3 cm in depth with three morsels of food (a small piece of peanut butter flavored cereal; Nestle, Columbia, MD). During the test, each mouse was given a 5-min period to find the buried food. The latency for the mouse to extract the buried food was recorded. The person performing the test and analyzing the data was blinded as to the genotype of the mice.

### 2.7 Elevated plus maze

Anxiety-like behavior was measured using the elevated plus maze [30]. The apparatus consists of two closed arms (30 × 5 × 15 cm) with high walls and two open arms (30 × 5 × 2.5 cm) with low walls. Each mouse was placed in the central area of the maze facing one of the open arms. Time spent in the open arms was measured for 5 min with the EthoVisionXT video-imaging system.

### 2.8 Open field test

For the open field test, animals were placed in the center of a defined open field region (43 cm × 43 cm) (Med Associates, Georgia, VT, USA) and left without disruption for 20 min. The center zone was defined as a 10.2 cm^2^ equidistant from the peripheral walls. The tracking software (Activity monitor version 4, Med-Associates) recorded the exploratory behavior. The apparatus was cleaned with 70% ethanol before testing the next mouse.

### 2.9 Electrophysiology

Hippocampal slices were prepared as described previously [30]. Briefly, the mice were killed using isoflurane gas, the brains were rapidly removed, and transverse slices were cut at a thickness of 350 μm. The slices were allowed to recover for at least 1 h in artificial cerebrospinal fluid (ACSF) at room temperature (RT) before recording. ACSF consisted of 120 mM NaCl, 2.5 mM KCl, 1.25 mM NaH2PO4, 26 mM NaHCO3, 1.3 mM MgSO4, 2.5 mM CaCl2, and 10 mM glucose (pH 7.4). The osmolarity of the ACSF was adjusted to 290 milliosmoles using a 5600 Vapor Pressure Osmometer (Wescor, Inc.). Stimuli (30 ms every 20 s) were delivered with a fine bipolar tungsten electrode to activate Schaffer collateral/commissural afferents. LTP was induced with a train of titanic stimulation (100 Hz for 1 s). All recordings were performed at 30–32°C. Data were collected using a MultiClamp 700B amplifier (Molecular Devices). Signals were filtered at 2 kHz, digitized at 10 kHz with a Digidata 1440A Data Acquisition System, and analyzed using pCLAMP 10 software (Molecular Devices).

### 2.10 Microarray

Gene expression analysis was performed on hippocampal tissue mRNA samples from APP/PS1 and WT littermates. RNA was purified with the Nucleospin RNA isolation kit (catalog no. 740955.250; Macherey-Nagel) with initial quantitation conducted using a NanoDrop ND-1000 spectrophotometer (Thermo Fisher Scientific). The quality of the RNA was inspected using a 2100 Bioanalyzer (Agilent Technologies). The microarray was performed by the Gene Expression and Genomics core facility (NIA) and analyzed with JMP, a SAS statistical Suite, combined with R. The initial fluorescent signals were first extracted from the Agilent result files with the background signal testing p values to ensure the validation of the signals. Then the data was transformed by log2 function and normalized with quantile normalization. Before doing further analysis, the data was standardized by z-transformation to fit for modeling. Then the samples went through the quality evaluation by sample level clustering, sample correlation with scatter plots, and principal component analysis to filter out outliers to ensure the analysis quality. The filtered samples were then organized according to their study groups to perform pairwise statistical analysis and variance test. The z-test with multiple comparator correction was adopted, along with one-way ANOVA variance test. The probes with ANOVA p value ≥ 0.05 and background signal ranking variance test p value ≥ 0.01 were filtered out in the whole set. For each pairwise comparison, we calculated z-test p value and zratio between pairs. We keep the probes for each pairwise comparison with all of the cutoffs: (1) average signal normalized zcore ≥ 0, (2) z-test p ≤ 0.05 (3) fdr ≤ 0.30 (we can adjust this according to different need) (4) |zratio| ≥ 1.5 in the subset probes of our global AVOVA and background filtering. The probes were further averaged on genes for the statistically significant gene list for each pair comparison with their zratios. Overrepresentation analysis of the genes which had an FDR of ≤ 0.05 were subjected to KEGG analysis for each pairwise analysis. We required at least 3 genes in each reported significant term in the data set. All the genes in the respective significant terms were collected.

### 2.11 Western blots

Mouse brain tissues were homogenized in 1× RIPA lysis buffer (Cell Signaling, #9806S) containing protease inhibitors cocktail, and halt phosphatase inhibitor cocktail (Roche, Indianapolis, IN, USA). Collected samples were sonicated on ice and centrifuged at 10,000 × g for 10 min at 4°C. The protein concentration was determined with Bradford reagent. 15 μg proteins were separated on 4-15% Bis-Tris gel (Bio-Rad Laboratories, #5671085) and transferred to PVDF blotting membranes. Membranes were then blocked for 1h at RT in TBS-T (500 mM NaCl, 20 mM Tris, 0.1% Tween 20) supplemented with 5% non-fat dried milk. Subsequently membranes were incubated overnight at 4°C with primary antibodies followed by 1 h at RT with HRP-conjugated secondary antibodies. Proteins were detected using an enhanced chemiluminescent detection system (EMD Millipore, #WBKLS0500) and ChemiDoc Imaging System (Bio-Rad Laboratories, #12003153, CA, USA). Quantification was performed using ImageJ. Antibodies used were: Phospho-Tau (Thr231) (catalog no. MN1040; ThermoFisher); Phospho-Tau (Thr181) (catalog no. MN1050; ThermoFisher); Phospho-Tau (Ser202, Thr205) (catalog no. MN1020; ThermoFisher); Tau antibody (catalog no. MN1000; ThermoFisher); Phospho-NF-κB p65 (rabbit) (catalog no. 3033; Cell signaling); GFAP antibody (catalog no. Z0334; 1:2000; Dako); Sirt3 antibody (catalog no. 2627; Cell signaling); Pink1 (catalog no. 23274-1-AP; Proteintech); Parkin (catalog no. 4211; Cell signaling); NLRP3 (catalog no. 15101; 1:500; Cell signaling); AIM2 (catalog no. 66902-1-Ig; Proteintech); STAT3 (catalog no. 10253-2-AP; Proteintech); p-STAT3 (catalog no. 9145; Cell Signaling); Bax (catalog no. 2722; 1:2000; Cell signaling); γH2AX (catalog no. 9718; Cell signaling); Ctsw (catalog no. LS-C712097-200; LSBio); Ctsd (catalog no. 21327-1-AP; Proteintech); Ctsz (catalog no. 16578-1-AP; Proteintech); β-Actin (catalog no. A2228; 1:5000; Sigma-Aldrich). Secondary antibodies including anti-mouse immunoglobulin G (IgG; 1010-05) and anti-rabbit IgG (catalog no. 4010-05) were obtained from Southern biotech. Primary antibodies were used at a 1:1000 dilution (unless otherwise stated), with secondary antibodies used at a 1:5000 dilution (unless otherwise stated). All antibodies were validated for use in mouse/human tissues based on previous publications. Detailed antibody validation profiles are available on the websites of the companies the antibodies were sourced from.

Membranes were then blocked for 1 h at RT in TBST (500 mM NaCl, 20 mM Tris, 0.1% Tween 20) supplemented with 5% non-fat dried milk and bovine serum albumin (BSA). Subsequently membranes were incubated overnight at 4°C with primary antibodies followed by 1 h at RT with HRP-conjugated secondary antibodies. Proteins were detected using an enhanced chemiluminescent detection system (EMD Millipore, catalog no. WBKLS0500) and ChemiDoc Imaging System (Bio-Rad Laboratories, catalog no. 12003153, CA, USA).

### 2.12 Mitochondrial purification

Mitochondrial extraction was adapted and modified from previous studies [32]. HMC3 cells with or without Aβ (10 µM) or UA (20 µM) treatment were washed with 1×PBS and harvested in mitochondria isolation buffer (20 mM HEPES, pH 7.5, 1.5 mM MgCl_2_, 1 mM EDTA, 1 mM EGTA, 210 mM sucrose, and 70 mM mannitol). Cell suspension was subjected to 60 gentle strokes using Dounce homogenizer. The supernatant obtained was again centrifuged at 10,000 g for 15 minutes at 4°C. The resultant mitochondrial pellet was washed twice and suspended in 200 μl buffer containing 250 mM sucrose, 5 mM magnesium acetate, 40 mM potassium acetate, 10 mM sodium succinate, 1 mM DTT, and 20 mM HEPES/KOH, pH 7.4.

### 2.13 Enzyme-linked immunosorbent assay (ELISA) for Aβ

Mouse hippocampal and PFC tissue were homogenized with RIPA buffer containing protease inhibitors as previously reported [30]. The levels of soluble and insoluble human Aβ_40_ and Aβ_42_ in these extracts were quantified using ELISA kits (Thermo Fisher Scientific, catalog no. KHB3442 for Aβ_40_ and KHB3482 for Aβ_42_) according to the manufacturer’s protocols. The HMC3 cell and 293 APP-Swedish cell supernatants were analyzed by ELISA according to manufacturer’s instructions. Briefly, cell supernatants were collected in centrifuges, after centrifugation 5 min at 10000 g, samples were added into microplate well and incubated with a fixed amount of target on the solid phase supporter, then added the biotinylated detection antibody specific to the target. Next, Adivin-HRP was added to each microplate well and incubated. After the TMB solution was added to each well, the enzyme-substrate reaction was terminated by the addition of a sulfuric acid solution, and the ODs was measured at a wavelength of 450 nm. Human IL-1β was measured by IL-1β kit (Thermo Fisher, catalog no. 88-7261). Human Aβ_42_ was measured by Aβ42 kit (ExCell Bio, catalog no. 22A23901).

### 2.14 RNA extraction and quantitative real-time PCR

RNA was extracted from mouse brain tissues as reported previously [14]. cDNA was synthesized using PrimeScript^TM^ RT reagent kit (Takara, catalog no. RR037A), and qPCR analysis was done with power SYBR Green PCR master mix (Thermo Fisher, catalog no. A46109). The primers used to amplify each transcript were as following: *Ctsz* (Forward: 5’-GGC CAG ACT TGC TAC CAT CC-3’ and Reverse: 5’-ACA CCG TTC ACA TTT CTC CAG-3’), *IL-1β* (Forward: 5’-GCA ACT GTT CCT GAA CTC AAC T-3’ and Reverse: 5’-ATC TTT TGG GGT CCG TCA ACT-3’), *IL-6* (Forward: 5’-GCC CAG CTA TGA ACT CCT TCT-3’ and Reverse: 5’-GAA GGC AGC AGG CAA CAC-3’), *Lamp1* (Forward: 5’-AGG CCA CTG TGG GAA ACT CAT ACA-3’ and Reverse: 5’-TTC CAC AGA CCC AAA CCT GTC ACT-3’), *IL-10* (Forward: 5’-CGG GAA GAC AAT AAC TGC ACC C-3’ and Reverse: 5’-CGGTTAGCAGTATGTTGTCCAGC-3’), *Trem2* (Forward: 5’-CTG GAA CCG TCA CCA TCA CTC-3’ and Reverse: 5’-CGA AAC TCG ATG ACT CCT CGG-3’); *Tyrobp* (Forward: 5’-GTG ACT TGG TGT TGA CTC TGC TG-3’ and Reverse: 5’-GAT AAG GCG ACT CAG TCT CAG C-3’), *Ctsh* (Forward: 5’-ACC GTG AAC GCC ATA GAA AAG-3’ and Reverse: 5’-TGA GCA ATT CTG AGG CTC TGA-3’), *Ctsa* (Forward: 5’-CCC TCT TTC CGG CAA TAC TCC-3’ and Reverse: 5’-CGG GGC TGT TCT TTG GGT C-3’), *Gapdh* (Forward: 5’-AGG TCG GTG TGA ACG GAT TTG-3’ and Reverse: 5’-GGG GTC GTT GAT GGC AAC A-3’).

### 2.15 Gene expression analysis by NanoString Technologies

NanoString analysis was performed on the hippocampus of AD and ADP mice with Veh or UA treatment and their wild type (WT) littermates. Total RNA was purified with a PureLink™ RNA Mini Kit (Thermo Fisher Scientific, catalog no. 12183018A) as per the manufacturer’s protocol. Purified RNA was quantified on a NanoDrop ND-1000 spectrophotometer and diluted in nuclease-free water to 20 ng/uL. It was hybridized in CodeSet Master mix carrying hybridization buffer, Reporter Code Set, and Capture Probe Set for 16 to 24 hours at 65°C (NanoString Technologies, MAN-10056-05) and then applied to the nCounter Prep Station. The Prep Station can process up to 12 samples per run in approximately 2.5 to 3 hours depending on which protocol is used. We loaded the hybridized RNA onto the nCounter Prep Station for immobilization in the sample cartridge according to the manufacturer’s high sensitivity protocol (MAN-C0035). The sample cartridge was subsequently processed for 2.5 hours in the nCounter Analysis System. Next, the nCounter Digital Analyzer which was a multi-channel epifluorescence scanner collected data by taking images of the immobilized fluorescent reporters in the sample cartridge with a CCD camera through a microscope objective lens. The results were directly downloaded from the digital analyzer in RCC files format. NanoString Advanced analysis (nSolver 4.0) was used for data analysis. Genes with a fold-change cut-off of ≥ |1.5| and p-value less than 0.05 were considered statistically significant. Pathway terms were considered significant if they had a Gene Set Enrichment score of 1.2.

The NanoString mouse AD panel was analyzed nSolver Advanced analysis and by ROSALIND® (https://rosalind.bio/), with a HyperScale architecture developed by ROSALIND, Inc. (San Diego, CA). Read distribution percentages, violin plots, identity heatmaps, and sample MDS plots were generated as part of the QC step. Normalization, fold changes and p-values were calculated using criteria provided by NanoString. ROSALIND® follows the nCounter® Advanced Analysis protocol of dividing counts within a lane by the geometric mean of the normalizer probes from the same lane. Housekeeping probes to be used for normalization are selected based on the geNorm algorithm as implemented in the NormqPCR R library [33]. Fold changes and pValues are calculated using the fast method as described in the nCounter® Advanced Analysis 2.0 User Manual. Venn diagrams were created by Venny 2.1 (https://bioinfogp.cnb.csic.es/tools/venny/). Clustering of genes for heatmaps of differentially expressed genes was done using http://www.heatmapper.ca/expression/. Enrichment was calculated relative to a set of background genes relevant for the experiment, ≥ 1.2 was considered significant.

### 2.16 Ctsz/Ctsw ELISA assay

Ctsz/Ctsw assays were measured using a mouse ELISA kit (MyBioSource, catalog no. MBS455784 and MBS7218953, San Diego, CA, USA) following the manufacturer’s protocol. This assay is based on Ctsz/Ctsw antibody-Ctsz/Ctsw antigen interactions and an HRP colorimetric detection system to detect Ctsz/Ctsw antigen targets in samples. The hippocampus tissue was homogenized with PBS (pH 7.0 - 7.2) on ice and centrifuged at 10,000 × g for 20 min at 4°C. The supernatants were collected and transferred to 96-well clear plate which is pre-coated with an antibody specific to Ctsz/Ctsw with a flat bottom. Brain samples were incubated for total 4 hours at 37°C and washed twice with 300 μL of 1× wash solution to each well using a multi-channel pipette for a total of 10 times. Substrate solution to each well subsequently added and incubated for 15 – 20 min at 37°C after covering with foil or plate sealer. The enzyme-substrate reaction was terminated by the addition of sulphuric acid (stop solution) and the absorbance value of each well was measured at a wavelength of 450 nm on microplate ELISA reader (Microplate Manager v5.2.1 software, Bio-Rad Laboratories).

### 2.17 Cytokine assay

Mouse eye bleeds were collected in EDTA-treated tubes. After centrifugation, the supernatant was flash frozen. Plasma diluted 1:2 was used to detect cytokines and chemokines by use of the 31-Plex Cytokine/Chemokine array (Eve Technologies).

### 2.18 Preparation of Aβ peptides

The peptides were prepared according to the protocols described previously [14]. Briefly, hexafluoroisopropanol (HFIP)-treated Aβ_42_ peptides (Chinese peptide) were resuspended in DMSO, then diluted to a concentration of 100 µM with DMEM/F12 and incubated at 4°C for 24 h. After centrifugation 10 min at 14,000 g, the supernatant with soluble Aβ_42_ was added to culture cells.

### 2.19 DqBSA assay

HMC3 cells were cultured in Minimum Essential Medium Eagle (Merck) containing 10% fetal bovine serum and 1% GlutaMAX (Thermo Fisher Sceintific) with or without 10 µM UA for 6 days. Cells were split and the following day dqBSA (Invitrogen, catalog no. D12051) was added in HBSS with or without 10 µM UA or bafilomycin A1 (Tocris Bioscience). dqBSA breakdown was imaged every hour using the Incucyte® Zoom live cell analysis system. A basic analysis using the Incucyte® Zoom software was conducted to score the total red object integrated intensity (RCU x µm^2^/image) over time. The linear range was used to determine the rate of degradation as done previously (https://www.ncbi.nlm.nih.gov/pmc/articles/PMC6693470/).

### 2.20 Measurement of lysosomal pH

The LysoSensor™ Yellow/Blue DND-160 Kit (Yeasen, #40768ES50) was used according to the manufacturer’s instructions. Briefly, HMC3 cells were seeded into glass bottom cell culture dish, and were transfected with (20 nM) siRNAs, 4 h later were cocultured with 10 μM Aβ_42_, 30 μM UA and 3 μM CTSZ Inhibitor or together for 48 h, diluting the drug stocks in Eagle’s Minimum Essential Medium with 2% fetal bovine serum, and loaded with 3 μM LysoSensor Yellow/Blue DND-160 of MEM for 30-40 min at 37°C. Then, the cells were washed 3 times with PBS buffer and immediately observed by fluorescence microscopy (OLYMPUS).

### 2.21 Randomization and blinding

Animal/samples (mice) were assigned randomly to the various experimental groups, and mice were randomly selected for the behavioral experiments. In data collection and analysis (for example, mouse behavioral studies, mouse imaging data analysis), the performer(s) was (were) blinded to the experimental design.

### 2.22 Statistical analysis

Prism 9.0 (GraphPad Software) was used for the statistical analysis. Data shown are the mean ± SEM with *P* ≤ 0.05 considered statistically significant. Two-tailed unpaired t-tests were used for comparisons between two groups. Group differences were analyzed with one-way analysis of variance (ANOVA) followed by Tukey’s multiple comparisons test or two-way ANOVA followed by Tukey’s multiple comparisons test for multiple groups. No statistical methods were used to predetermine sample sizes, but our samples sizes (mouse experiments) were similar to those reported in previous publications.

### 2.23 Data availability

The microarray GEO accession number for the data reported in this paper is GSE212972, GSE214406 and GSE214416. All data are available from the corresponding author upon reasonable request.

## 3 RESULTS

### 3.1 Early and long-term UA treatment improves learning and memory in APP/PS1 mice

APP/PS1 mice are widely used in AD research and they develop Aβ pathology and behavioral deficits by 6 months [21, 34, 35]. Thus, we sought to initiate UA treatment before significant AD phenotypes occur. Notably, mitophagy defects may occur before the main pathologies of AD, thus early intervention testing with mitophagy stimulation is important. Two-month-old APP/PS1 mice were exposed to UA for 5 months and then behavioral tests were performed, and mouse brains were collected (Fig. 1a). In the Morris water maze experiment, we found that APP/PS1 mice took more time to reach the hidden platform than WT mice, indicating that their learning ability had decreased (Fig. 1b). Compared with APP/PS1 mice treated with vehicle, APP/PS1 mice treated with UA took significantly less time to reach the platform, like that of WT (Fig. 1b). This result shows that UA administered for 5 months significantly improved the learning ability of APP/PS1 mice. During the experiment, the swimming speed of mice in each group was similar (Fig. 1c), demonstrating that UA improved learning ability independent of motor function. In addition, we also used another cognitive function test, Y-maze, to evaluate the long-term UA treatment effect on memory. 5-month UA treatment significantly improved the memory ability of APP/PS1 mice (Fig. 1d).

**Figure 1.**
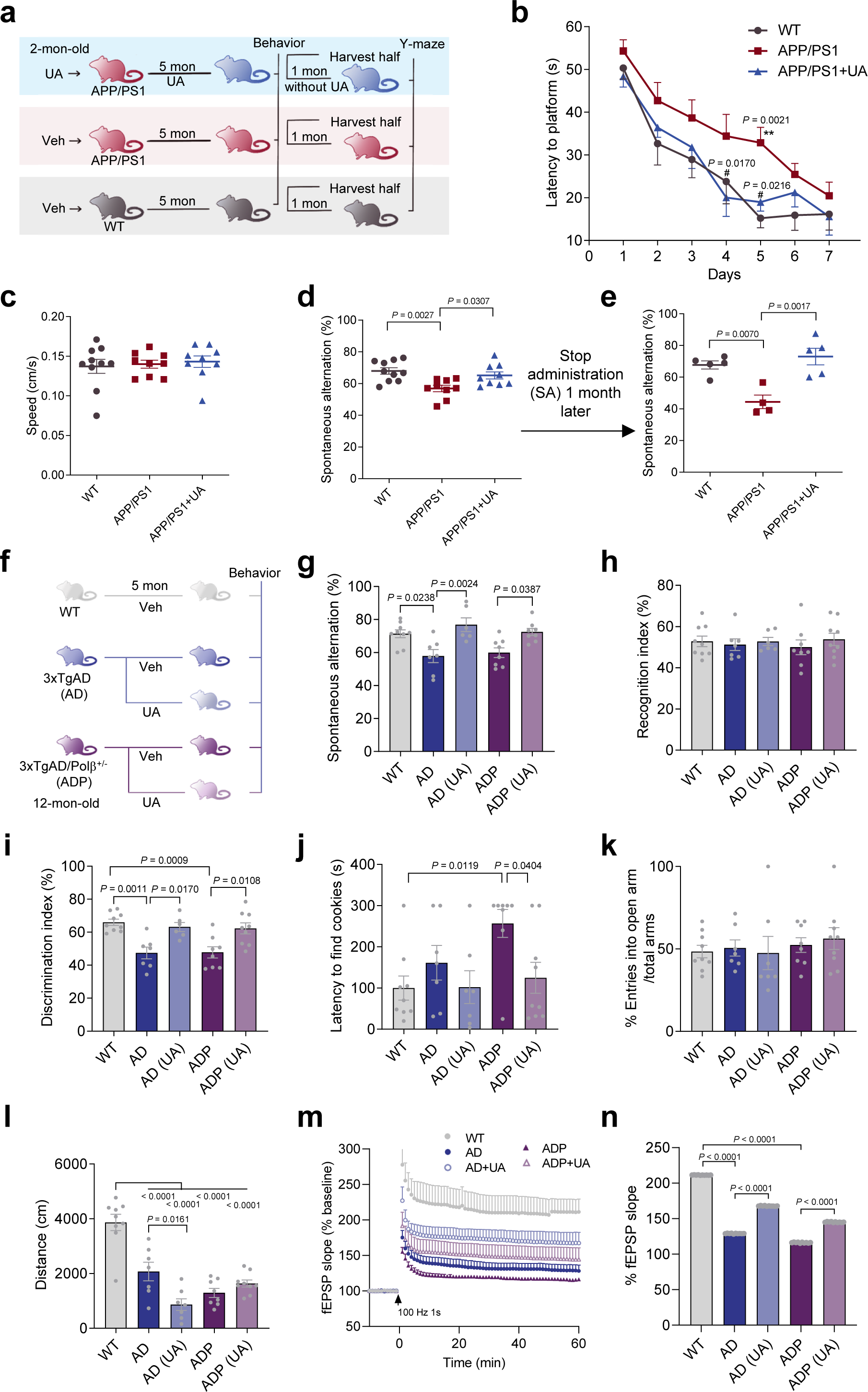
UA improved learning and memory ability in APP/PS1, 3xTgAD (AD) and 3xTgAD/Polβ^+/-^ (ADP) mice. (**a**) Experimental design for UA treated APP/PS1 mice. (**b**) The learning phase of Morris water maze. The latency to find the hidden platform of WT, APP/PS1 and UA-treated APP/PS1 mice. \*\**P* < 0.01, APP/PS1 mice compared to WT mice. #*P* < 0.01, UA-treated APP/PS1 mice compared to vehicle-treated APP/PS1 mice; For learning curve comparison: *P* = 0.002, APP/PS1 compared to WT; *P* = 0.0021, APP/PS1+UA compared to APP/PS1, by Pearson’s correlation. n = 9-10 male mice. (**c**) The swimming speed in Morris water maze, n = 9-10 male mice. (**d**) Y-maze test of 5-month UA-treated APP/PS1, vehicle-treated APP/PS1 mice and WT mice, n = 9-10 male mice. (**e**) Y-maze test of 5-month UA-treated APP/PS1, vehicle-treated APP/PS1 mice and WT mice, after stop administration of UA for 1 month, n = 4-5 male mice. (**f**) Experimental design for UA treated AD and ADP mice. (**g**) Y-maze test of 5-month UA-treated or vehicle-treated AD, ADP and vehicle-treated WT mice, n = 6-9 mice. (**h-i**) Object recognition test of 5-month UA-treated or vehicle-treated AD, ADP and vehicle-treated WT mice, n = 6-9 mice. (**j**) The latency to find cookies in buried food test of the mice, n = 7-9 mice. (**k**) Elevated plus maze of the mice in (B), n = 7-9 mice. (**l**) Open field test of the mice. (**m-n**) Results of analyses of LTP measured at the Schaffer collateral synapses. Values are the mean and SEM of determinations made on 5-7 hippocampal slices from at least 5 different mice. EPSP, excitatory postsynaptic potential. Data were analyzed by two-way ANOVA with Tukey’s multiple comparisons test and Pearson’s correlation (**b**) or one-way ANOVA with Tukey’s multiple comparisons test (**c-e, g-l, n**). Data were shown as mean ± SEM. The APP/PS1 mice are all males. For g-i, WT (7 M and 2 F), AD (3 M and 4 F), AD + UA (3 M and 3 F), ADP (5 M and 3 F), ADP + UA (3 M and 5 F); for j-l, WT (7 M and 2 F), AD (3 M and 4 F), AD + UA (3 M and 4 F), ADP (5 M and 3 F), ADP + UA (4 M and 5 F). F, female, M, male.

We wondered if UA treatment would have a lasting impact on disease endpoints, so we measured Y-maze one month after discontinuing the 5 month-administration of UA. After one month without UA, APP/PS1 mice treated with UA still showed improved memory ability compared to APP/PS1 mice treated with vehicle (Fig. 1e). Motor function and anxiety were not changed in the APP/PS1 mice compared to WT mice, nor after UA treatment (supplementary Fig. 1a-d). These results show that 2-month-old APP/PS1 mice given 5-months of UA significantly improved learning and memory function in APP/PS1 mice, and this effect persisted one month after UA was discontinued.

### 3.2 Long-term UA treatment improved learning and memory in 3xTgAD and 3xTgAD/Polβ^+/-^ mice

To verify the therapeutic effects of early and long-term UA treatment on AD, we used two other AD models, the widely used AD and the DNA repair deficient ADP mice, which we developed [29]. The 3xTgAD mice are APP and tau transgenic mice, and we crossed them with DNA repair-deficient Polβ^+/-^ mice [29] and reported that they had many more human AD features than the 3xTgAD mice [29]. Decreased DNA repair activity has been found in AD patients, and we reported a base excision DNA repair defect at the level of DNA Polβ in human early postmortem AD brains [36]. In our AD and ADP mice, we typically find Aβ and tau pathology starting around 12 months [29]. For the AD and ADP mouse models, UA was administered orally from 12 months to 17 months (duration 5 months, Fig. 1f). UA significantly improved the memory of AD and ADP mice in the Y-maze test (Fig. 1g). In the object recognition test, the cognitive ability of AD and ADP mice to recognize two identical objects was like WT mice (Fig. 1h), but the ability to recognize novel objects had significantly decreased (Fig. 1i). After UA treatment, the novel object recognition ability of AD and ADP mice had significantly improved (Fig. 1i), showing that UA improved the cognitive ability of AD and ADP mice.

Sensory perception is altered in AD patients. AD patients commonly have olfactory impairment [37] and we have previously reported that our ADP mice have olfactory dysfunction [38, 39]. Thus, we tested the mice using the buried food smelling test. The latency of ADP mice to find the buried cookies was significantly longer than that of WT and long-term treatment with UA improved the olfactory function in these mice, while the AD mice did not have a significant smelling impairment (Fig. 1j).

To assess UA effects on anxiety, we first used the elevated plus maze test. AD and ADP mice had no anxiety-related behavior in this experiment and were not significantly different from WT (Fig. 1k). However, in the open field test, the walking distance of the AD and ADP groups were reduced relative to WT mice, and the walking distance of the UA-treated AD group was shorter than that of AD group, however, this effect was not observed in the ADP mice group (Fig. 1l and supplementary Fig. 1e-f). To further compare the effects of UA on the behavior of AD model mice by sexes, we divided female and male for statistical analysis. The results showed that female and male mice had similar trends (supplementary Fig. 1e-f, right panels; and supplementary Fig. 2).

The decline of cognitive function in AD patients is related to the loss of synaptic function. Therefore, we studied long-term potentiation in the mice using electrophysiological experiments. The results showed that UA significantly improved long-term potentiation of AD and ADP mice (Fig. 1m-n). Collectively, these results indicate that long-term administration of UA improves the cognitive, olfactory, and synaptic functions of neurons in AD and ADP mice.

### 3.3 UA decreased Aβ accumulation and Tau phosphorylation in AD mice

To investigate the effects of UA treatment on Aβ and phosphorylated Tau (pTau), we used APP/PS1, AD and ADP mice. In APP/PS1 mice, we performed ELISAs to examine whether UA treatment decreased the levels of Aβ in the hippocampus and prefrontal cortex (PFC), known as two of the earliest and most affected brain areas in AD [40]. There were no significant changes in insoluble and soluble of Aβ_42_ and Aβ_40_ in the hippocampus of APP/PS1 mice after long-term UA treatment (Fig. 2a-d). However, in the PFC, both the soluble and insoluble Aβ_42_ were significantly decreased after long-term UA treatment (Fig. 2a and 2c). The levels of soluble Aβ_40_ were also reduced in the PFC of APP/PS1 mice (Fig. 3d), while insoluble Aβ_40_ showed no changes (Fig. 2b).

**Figure 2.**
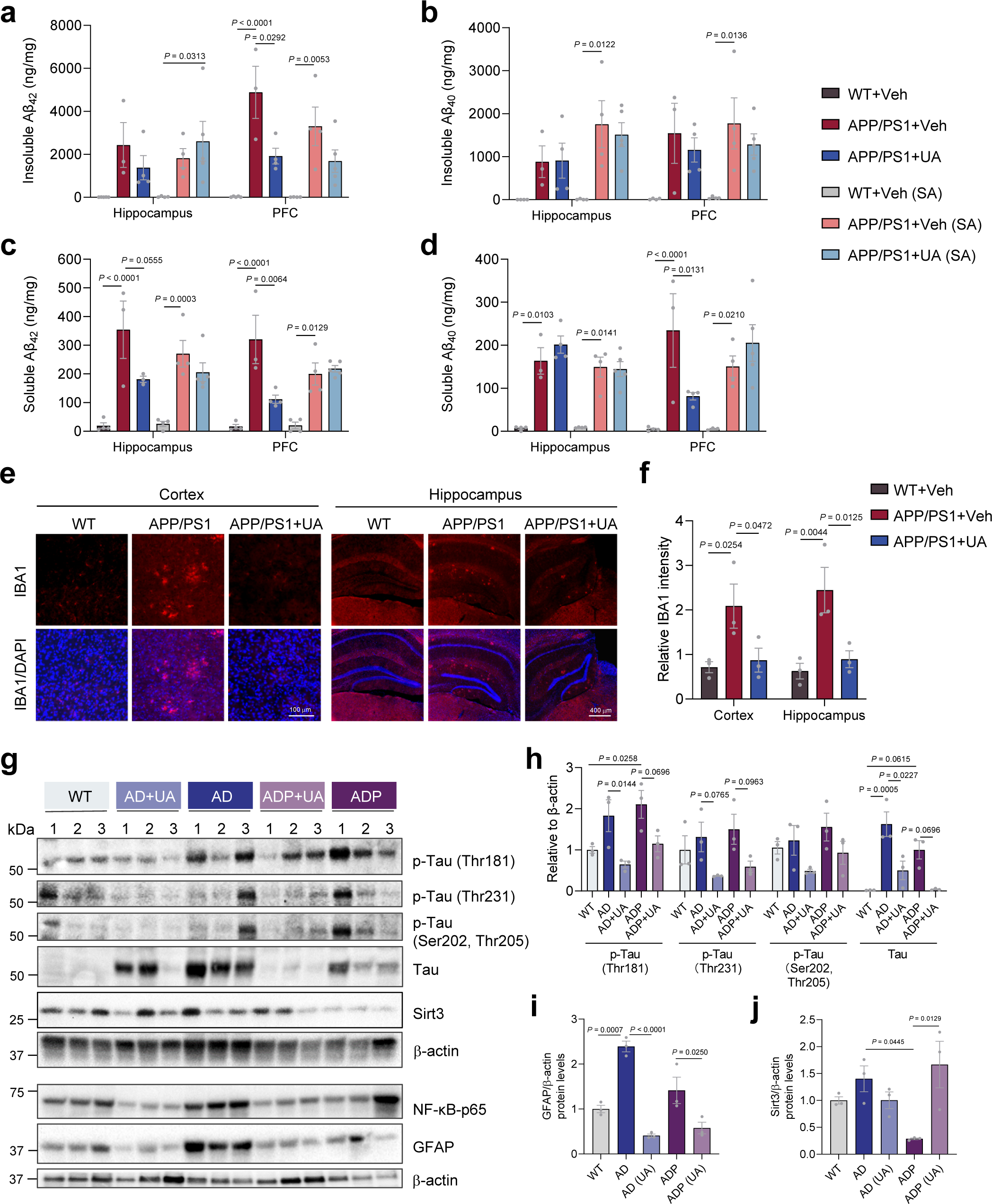
UA decreased Aβ accumulation and Tau phosphorylation in AD mice. (**a-d**) The effects of UA treatment on Aβ productions in both hippocampus and PFC. Insoluble Aβ_42_ (a), insoluble Aβ_40_ (b), soluble Aβ_42_ (c), and soluble Aβ_40_ (d) levels in all groups. Data were shown as Mean ± SEM (n = 3 male mice in the AD group; n = 5 male mice in the AD+UA-washout group; n = 4 male mice in all the other groups; \**P* < 0.05; two-way ANOVA with Tukey’s multiple comparisons test). (**e-f**) The representative images and quantification of immunostaining of IBA1 in the cortex and hippocampus in WT, vehicle- or UA-treated APP/PS1 mice. n = 3 male mice per group. Scale bars, 100 μm and 400 μm, as indicated. (**g**) Western blot data showing the effects of UA treatment on the expression level of proteins involved in Aβ accumulation and Tau phosphorylation in the hippocampus of WT, 3xAD, and 3xAD/Polβ^+/-^ mice, n = 3 mice per group (**h**) Quantification of protein levels for p-Tau (Thr181), p-Tau (Thr231), p-Tau (Ser202, Thr205), and total Tau. (**i-j**) Quantification of protein levels for GFAP and Sirt3. For h-j, n = 3 mice per group. All samples were normalized to respective loading controls β-actin. Data were analyzed by one-way ANOVA with Tukey’s multiple comparisons test (**i-j**) or two-way ANOVA with Tukey’s multiple comparisons test (**a-d**, **f**, **h**). Data were shown as Mean ± SEM. The APP/PS1 mice are all males. For AD, AD+UA: 2 M and 1 F; For WT, ADP and ADP+UA, 1 M and 2 F. M, male; F, female.

**Figure 3.**
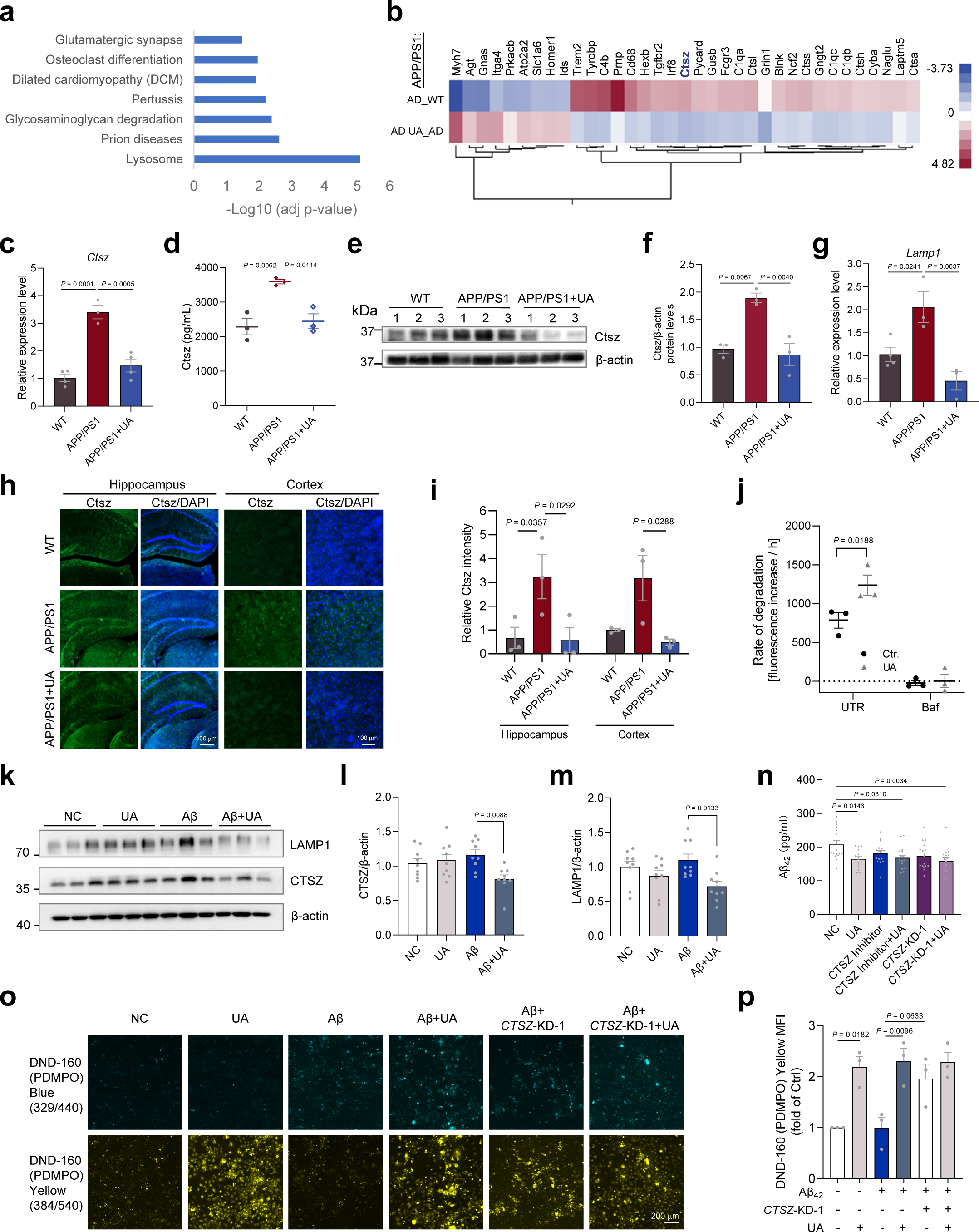
UA restored lysosomal functions through regulating Ctsz in APP/PS1 mice. (**a**) KEGG term analysis of gene expression microarray from the hippocampus of APP/PS1 mice. The most changed terms in AD compared to WT hippocampus were shown. (**b**) The FDR selected genes (cutoff 0.05) from each term in (a) from hippocampal microarray of APP/PS1 compared to WT and APP/PS1 compared to APP/PS1 with UA treatment are shown. (**c**) qPCR analysis for the relative gene expression of *Ctsz*. (**d**) Measurement of Ctsz levels by ELISA assay in hippocampus of APP/PS1 or WT mice with and without UA treatment. (**e-f**) Western blots of the Ctsz protein expression levels in hippocampus of APP/PS1 with or without UA treatment. (**g**) qPCR analysis for the relative genes expression of *Lamp1* in the hippocampus. For qPCR, n = 3-4 mice. (**h**) The representative images of immunostaining of Ctsz in the hippocampus and cortex of WT, vehicle- or UA-treated APP/PS1 mice. n = 3 mice per group. Scale bars, 400 μm and 100 μm, as indicated. (**i**) The quantification of Fig. 3h. (**j**) DqBSA degradation assay in HMC3 cells with UA treatment or Baf treatment. (**k-m**) Representative western blots showing LAMP1 and CTSZ and quantitation of protein levels in UA or Aβ or UA + Aβ treated HMC3 cells, n = 9 independent repeats. (**n**) ELISA for Aβ42 in UA or Ctsz inhibitor or si-*CTSZ* treated 293-APPsw cell supernatant. (**o-p**) The measurement of lysosomal pH by using the LysoSensor™ Yellow/Blue DND-160 Kit, HMC3 cells were treated with siRNAs, 10 μM Aβ_42_, 30 μM UA and 3 μM CTSZ Inhibitor or together for 48 h. Scale bars, 200 μm. Data were shown as mean ± SEM. Data were analyzed by one-way ANOVA with Tukey’s multiple comparisons test (**c-d**, **f-g**, **l-n**, **p**) or two-way ANOVA with Tukey’s multiple comparisons test (**i, j**). All APP/PS1 strain mice used are males.

One month after discontinuing UA administration, no significant changes were observed for Aβ_42_ and Aβ_40_ in the hippocampus of APP/PS1 mice (Fig. 2a, c). We also did not detect decreased levels of Aβ_40_ and Aβ_42_ in PFC in the group without UA administration for one month (Stop administration, SA), suggesting that the beneficial effects observed in the PFC vanished in APP/PS1 mice one month after stopping UA administration (Fig. 2a-d, SA group). The results show that UA treatment reduced Aβ accumulation in the PFC but not in the hippocampus of APP/PS1 mice, and that the effects disappeared if UA treatment was suspended for one month.

Abnormally activated neuroinflammation is an important pathological feature of AD. We detected activated microglia cells in the brains of AD mice. The number of IBA1-positive cells increased significantly in the cortex and hippocampus of APP/PS1 mice, and UA reduced these abnormally activated microglia cells dramatically in the cortex and hippocampus of APP/PS1 mice (Fig. 2e-f).

Next, we investigated the effects of UA treatment on Tau phosphorylation in ADP brains. Previous studies have shown that phosphorylation sites of Tau, including Thr181, Thr231, and Ser202/Thr205, were associated with disease progression [41]. Notably, UA treatment significantly decreased total Tau levels at Thr181 and Thr231 in both AD and ADP mice (Fig. 2g-h). UA treatment also decreased pTau levels in ADP but not in AD brains (Fig. 2g-h). Thus, our results showed that UA treatment suppressed the accumulation of total Tau and pTau in brains.

Glial fibrillary acidic protein (GFAP), known as a marker for astrocytes, is associated with AD pathology as an early marker of brain Aβ status [42]. We found that GFAP was remarkably increased in AD brains and normalized after UA treatment (Fig. 2g and 2i). However, there was no differences of the expression of GFAP between ADP and WT mice. It was reported that the neurotoxic Aβ peptides induced NF-κB activation in AD brains [43]. Since inflammation is a major cause of Aβ and pTau [44], we explored whether UA treatment attenuated the inflammatory reactions in AD mice. Interestingly, long-term UA treatment tended to decrease the expression level of activated NF-κB, as determined by increased p65 in the hippocampus of AD mice compared with WT mice (Fig. 2g and supplementary Fig. 3a). However, there was no difference between WT and AD/ADP mice without UA treatment, suggesting that the downregulated NF-kB specifically responded to UA treatment (Fig. 2g and supplementary Fig. 3a). In addition, sirtuin 3 (SIRT3) is a mitochondrial deacetylase, and its dysfunction is strongly associated with the pathogenesis of AD [45]. A previous study revealed that Aβ increased total tau levels through the regulation of SIRT3 in AD adults [46]. Interestingly, our results indicated that Sirt3 tended to decrease in the ADP mice but not in the AD mice (Fig. 2g and 2j). Moreover, the expression level of Sirt3 significantly increased after long-term UA treatment in ADP mice (Fig. 2j). Taken together, we speculate that UA might downregulate Aβ- and pTau-induced inflammation and DNA damage in AD brains.

### 3.4 UA restored lysosomal functions in part by regulating cathepsin Z

To further explore the underlying mechanisms related to UA treatment of AD, we performed gene expression analysis by microarray using hippocampal tissues from WT, untreated and treated APP/PS1 (AD) mice. KEGG terms showed that the most significantly altered term in UA-treated AD mice was related to lysosomes (Fig. 3a). The significantly differentially expressed genes are shown in Fig 3b. Cathepsin A (*Ctsa*), cathepsin H (*Ctsh*), cathepsin L (*Ctsl*), cathepsin S (*Ctss*), and cathepsin Z (*Ctsz*), were all significantly upregulated in AD brains and UA treatment decreased their expression (Fig. 3b). Cathepsins are vital lysosomal compartment enzymes responsible for intracellular protein degradation, energy metabolism, and immune responses [47]. Dysregulation of cathepsin proteases may contribute to the development of AD and Aβ productions [48]. Consistent with these results, our data showed a specific increase in cathepsin gene expression levels in AD brains, which might contribute to the progression of AD (Fig. 3b). The mRNA levels of the cathepsins (*Ctsa*, *Ctsh*, *Ctss*, *Ctsz*) were examined by quantitative real-time PCR (qPCR) in all brain samples (Fig. 3c, supplementary Fig. 3b-d). Among them, we found that *Ctsz* was significantly elevated in the hippocampus of AD brains, and UA normalized its expression to control levels (Fig. 3c). Ctsz is a member of the lysosomal cysteine cathepsin protease family, which comprises 11 members in humans [47]. In the context of neuroinflammation, Ctsz has been implicated in the development of inflammation, Interleukin-1β (IL-1β) production, and NLR Family Pyrin Domain Containing 3 (NLRP3)-inflammasome activation [49, 50]. We measured Ctsz levels to further evaluate the effects of UA in AD brains by ELISA. The Ctsz level was significantly higher in the hippocampus of APP/PS1 mice than in WT mice and decreased notably after UA treatment (Fig. 3d).

The mRNA expression levels of *Ctsa* and *Ctsh* were higher in the hippocampus of APP/PS1 mouse brains than controls, but UA did not affect them significantly (supplementary Fig. 3b-c). *Ctss* was not changed in the AD and UA-treated groups compared with WT (supplementary Fig. 3d). The protein levels of Ctsz were increased in AD mouse hippocampus and decreased after UA treatment (Fig. 3e and 3f). We additionally used immunostaining and found that Ctsz was higher in the cortex and hippocampus of the APP/PS1 mice and decreased in after UA treatment (Fig. 3h-i). The lysosomal marker, lysosomal-associated membrane protein 1 (*Lamp1*), was more highly expressed at the mRNA level in the hippocampus of AD mice and decreased significantly after UA treatment, suggesting that UA may contribute to down-regulating lysosomal degradation in AD (Fig. 3g).

To examine lysosomal degradation activity after UA treatment, we performed the Dye Quenched-Bovine Serum Albumin (DqBSA) assay and the lysosensor assay. In the DqBSA degradation assay, UA significantly increased DqBSA degradation in the HMC3 cells, and bafilomycin (Baf) was used as negative control (Fig. 3j), suggesting that UA increased lysosomal function.

Microglia are major immune cells of the central nervous system, and HMC3 human microglia were used for further study. HMC3 human microglia were treated with Aβ_42_ (10 μM) with or without UA for 48 h, and the protein levels of CTSZ and LAMP1 decreased when UA was added with Aβ_42_ (Fig. 3k-m). In addition, ELISA showed that UA decreased Aβ_42_ expression in the supernatant of HEK293-APPswe cells (Fig. 3n). Using the lysosensor kit to detect lysosomal pH, we found increased yellow DND160 signal in the cells treated with UA, suggesting that UA promotes acidification of lysosomes and thereby may improve lysosomal function (Fig. 3o). Note, Aβ-treatment reduced the cellular lysosome activity, and UA restored it. Interestingly, the UA-induced lysosomal activity was blocked after *CTSZ* knockdown, maybe because of *CTSZ* knockdown activates lysosome acidification by itself thereby rendering more difficult to see the stimulation of UA, or implicating that UA functions through CTSZ, at least by part, to improve the lysosome function (Fig. 3o-p). *CTSZ* knockdown efficiency is shown in supplementary Fig. 3e-f.

### 3.5 UA decreased inflammation in AD mice

CTSZ is associated with neurodegenerative diseases like AD and Huntington’s disease (HD) [51, 52] and is related to inflammation and Interleukin-1β (IL-1β) production [50]. We next examined the mRNA expression level of *IL-1β* and *IL-6* in the hippocampus to assess the effects of UA on inflammation. *IL-1β* was enhanced in AD mice and UA notably decreased its level (Fig. 4a). However, there was no difference in *IL-6* expression among all groups (Fig. 4b), suggesting that UA weakened inflammation mainly through the IL-1β-related signaling pathway. *IL-10* is an anti-inflammatory cytokine, which was decreased in AD mice and restored after UA treatment (Fig. 4c). Cytokine levels were also measured from cortex lysates using the Cytokine assay. We found that several pro-inflammatory cytokines were increased in AD mice and decreased after UA treatment, including IL-1α, Monocyte chemoattractant protein-1 (MCP-1), Macrophage Inflammatory Protein-1 Alpha (MIP-1α), tumor necrosis factor (TNFα), IL-2, and keratinocyte-derived cytokine (KC) (Fig. 4d-i). The IL-6 level did not show significant differences between the groups (supplementary Fig. 3g). Some other cytokines were changed, and most of them showed the same trends: increased in AD mice cortex lysates and decreased in UA-treated mice (supplementary Fig. 3h-s). To further investigate the mechanism by which UA enhances lysosome function and the correlation with CTSZ, HMC3 cells primed with LPS were treated with *CTSZ*-siRNA or CTSZ inhibitors, with or without Aβ_42_ and UA. In the Aβ_42_-treated condition, UA significantly reduced IL-1β expression in the HMC3 cell supernatant but did not reduce IL-1β in the *CTSZ*-knockdown. Consistent with the *CTSZ* knockdown results, CTSZ inhibitors also reduced IL-1β expression, and UA did not further reduce IL-1β in HMC3 cells treated with CTSZ inhibitors. These results suggest that UA reduces IL-1β in a way that is partly dependent on CTSZ (Fig. 4j).

**Figure 4.**
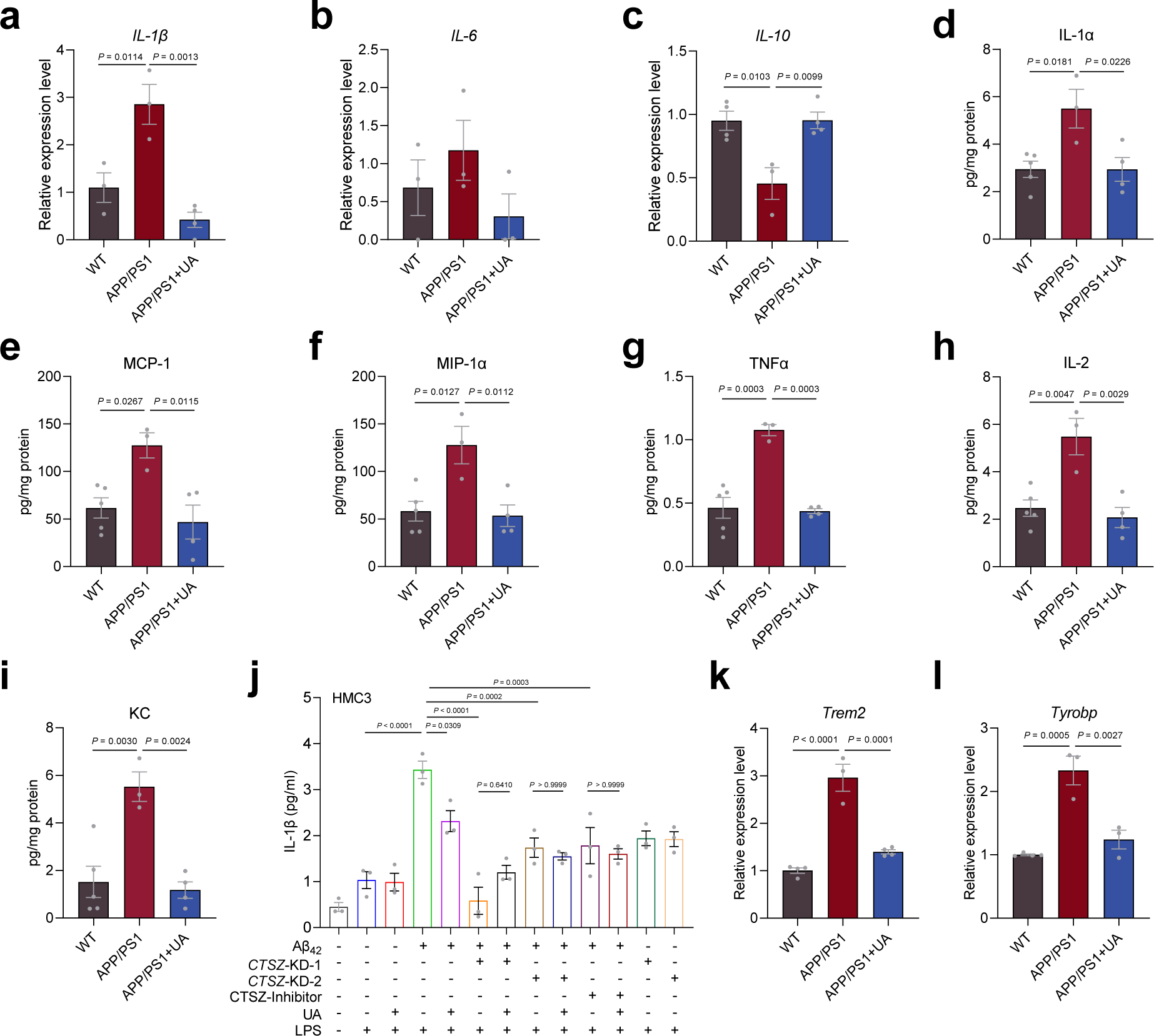
UA decreased pro-inflammatory factors in AD mice. (**a-c**) qPCR analysis for the relative genes expression of *IL-1β* (a), *IL-6* (b) and *IL-10* (c). (**d-i**) The cytokine assay of cytokine levels in mice cortex lysates, including IL-1α, MCP-1, MIP-1α, TNFα, IL-2, and KC, n = 5 in the WT group; n = 3 in the APP/PS1 group; n = 4 in the APP/PS1 + UA group. (**j**) ELISA of IL-1β in Aβ_42_ or UA or *Ctsz*-knockdown or Ctsz-inhibitor treated HMC3 cells. n = 3 biological duplicates. (**k-l**) qPCR analysis for the relative gene expression of *Trem2* (k), and *Tyrobp* (l). For qPCR, n = 3-4 mice. Data were shown as mean ± SEM. Data were analyzed by one-way ANOVA with Tukey’s multiple comparisons test (**a-l**). For APP/PS1 strain mice, all used are males.

The triggering receptor expressed on myeloid cell 2 (TREM2) binds to the tyrosine motif binding protein (TYROBP; also known as DAP12, DNAX-binding protein-12), a critical modulator in microglial biology and correlates with a high risk of AD development [53, 54]. Our RNA sequencing data by NanoString showed that the expression levels of *Trem2* and *Tyrobp* markedly increased in APP/PS1 mice and decreased dramatically after UA treatment in APP/PS1 brains (Fig. 3b), and this was verified by the mRNA expression results from the hippocampus tissues (Fig. 4k-l). CD68 is a lysosomal protein expressed at high levels by macrophages and activated microglia. We found that *CD68* was higher in the hippocampus of APP/PS1 mice and unchanged in UA-treated APP/PS1 mice (supplementary Fig. 4a). To investigate the effects of UA on neuroinflammation, we tested some inflammation-related proteins in mouse hippocampus and found that UA decreased NF-κB p105/p50, AIM2 and pSTAT3/STAT3 (supplementary Fig. 4b-f). The mTOR pathway has been implied in the regulation of autophagy and mitophagy [55]. We found that the mTOR expression level was decreased in APP/PS1 mouse brains and increased after UA treatment (supplementary Fig. 4g-h). UA also trended to decrease CD68 protein in APP/PS1 mice brains (supplementary Fig. 4g and 4i). Accordingly, these data suggest that UA administration reduces neuroinflammation in AD mice.

### 3.6 UA induced Sirtuin expression, mitophagy, and decreased DNA damage

The sirtuin family of proteins impacts both aging and mitophagy [56], thus we assessed the expression of Sirt1 and Sirt3 in the AD mouse cortex, and found that UA increased the expression of both proteins in APP/PS1 mice (Fig. 5a-c). In our previous study, we found that UA plays an important role in AD therapy as a mitophagy inducer [11]. Here, we also verified the effect of long-term administration of UA on mitophagy-related indicators in APP/PS1 mice. UA increased the expression of Parkin and BNIP3 (Fig. 5d and supplementary Fig. 5a and 5e), which were decreased in APP/PS1 mice, and tended to increase the expression of other mitophagy related proteins, such as Nix and Mfn2 (Fig. 5d and supplementary Fig. 5b-c). Long-term administration of UA was also found to dramatically reduce DNA damage responses, as assessed by γ-H2AX (Fig. 5d and f). We also investigated PARylation level and p-Parkin in UA-treated APP/PS1 mouse cortex samples, and found that UA decreased PARylation and p-Parkin in APP/PS1 mouse cortex samples, supporting that UA decreases DNA damage responses (supplementary Fig. 5d-f).

**Figure 5.**
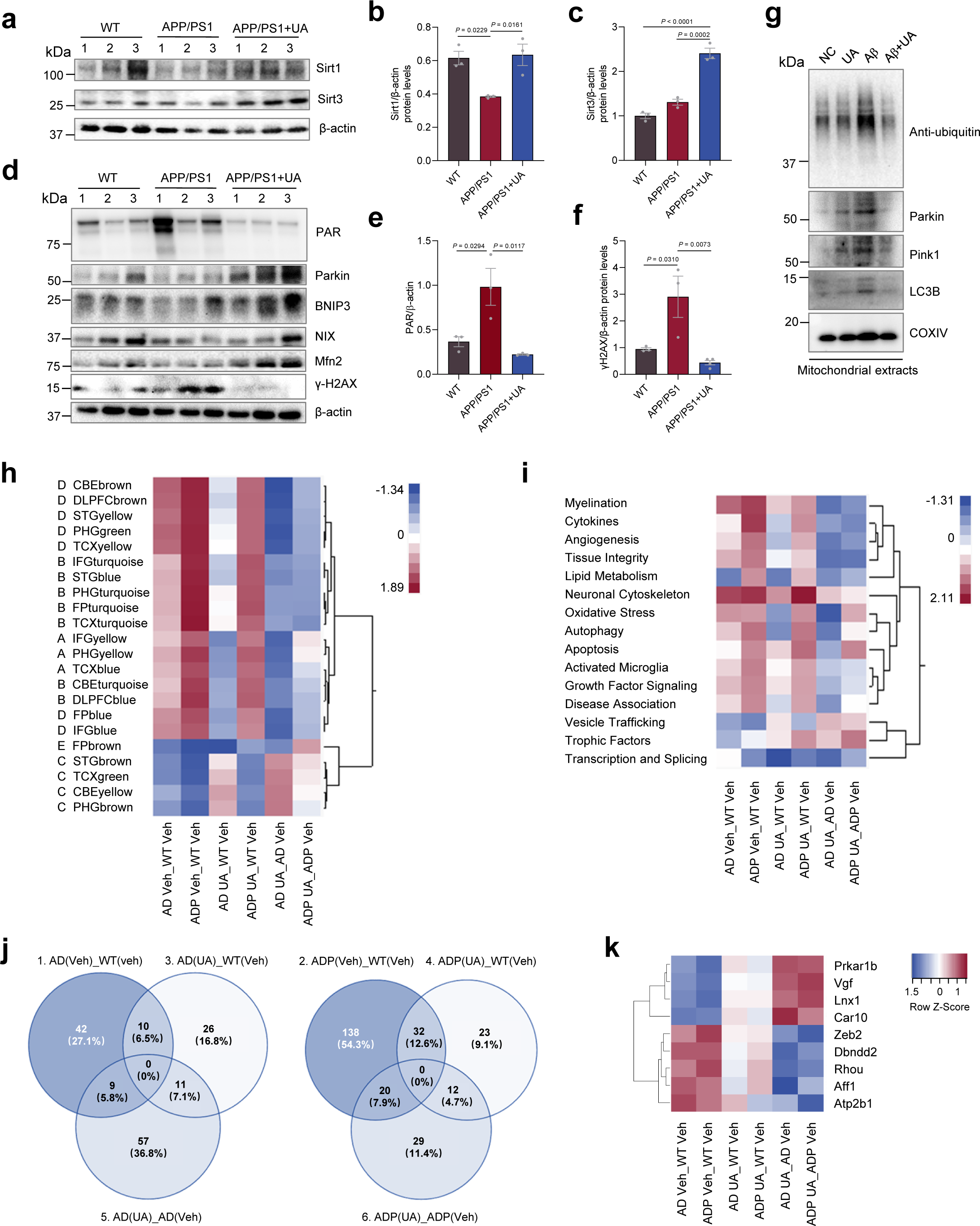
UA modulates both general immune and neuronal processes and AD-specific pathophysiological pathways. (**a-c**) Representative western blots showing Sirt1 and Sirt3 and quantitation of protein levels in the cortex of WT, APP/PS1, and APP/PS1 with UA mice, n = 3 male mice per group. (**d-f**) Representative western blots showing PAR, Parkin, BNIP3, NIX, Mfn2, γH2AX and quantitation of protein levels in the cortex of WT, APP/PS1, and APP/PS1 with UA mice, n = 3 male mice per group. (**g**) The representative western blots showing ubiquitin, Parkin, PINK1, LC3B and COXIV in the HMC3 cell of NC, UA, Aβ and Aβ + UA treatment, n = 3 independent repeats. (**h**) Heatmap in the hippocampus showing gene set enrichment analysis of significant terms from the various AD clusters defined by the NanoString mouse AD panel. Cluster designations A-E are shown to the left and defined below. Cluster A represents ECM organization; Cluster B, immune system; Cluster C, neuronal system; Cluster D, cell cycle, NMD; Cluster E, organelle biogenesis, cell stress response. (**i**) Heatmap in the in hippocampus showing gene set enrichment analysis of significant pathology terms defined by the NanoString mouse AD panel. (**j**) Venn diagram of AD and ADP mice showing significant genes (p-value ≤ 0.05) and their relative overlap between the treatment groups. (**k**) Set of significantly changed genes (fold-change ≥ |1.2| and *p*-value ≤ 0.05) changed similarly by UA in both AD and ADP. Data were shown as mean ± SEM. Data were analyzed by one-way ANOVA with Tukey’s multiple comparisons test (**b-c**, **e-f**).

Alterations in mitochondria and lysosomes are often both present in neurodegenerative diseases, suggesting a close relationship between mitochondria and lysosomes [57]. To investigate the effect of UA on mitophagy, we examined the markers of mitophagy in mitochondrial extracts. We used UA, Aβ, and Aβ + UA in HMC3 cells. Mitochondria were extracted, and proteins related to mitochondrial autophagy were assessed. Mitochondrial Parkin was significantly increased and ubiquitin, PINK1, and LC3B were tended to increase after Aβ treatment, and UA restored the expression of these mitophagy-related proteins to similar levels as the control group (Fig. 5g and supplementary Fig. 6a-d). The results indicate that acute Aβ stimulation leads to abnormal mitophagy-related protein expression in extracts, and UA restores these proteins to normal levels.

### 3.7 Targeting multiple mechanisms of pathology, UA modulates immune responses and AD-specific pathophysiological pathways

To assess gene expression changes after UA, we employed the NanoString Alzheimer’s Disease panel (Fig. 5h). The AD panel was designed to correlate key human disease processes and pathways with mRNA from mouse brains. It consists of genes associated with the primary molecular characteristics of AD [58]. The genes included 30 AD-associated brain region-specific gene co-expression modules defined by the Accelerating Medicines Partnership-Alzheimer’s Disease (AMP-AD) consortium and 23 neurodegenerative pathways and processes [58, 59]. The co-expression modules are further partitioned into five distinct consensus clusters that share minimal gene overlap (clusters A-E). Each cluster is associated with various concepts such as A, extracellular matrix organization; B, immune system; C, neuronal system; D, cell cycle and non-sense mediated decay; E, organelle biogenesis and cell cycle response. Consistent with the previous article, we set our Gene Set Enrichment Analysis threshold for significance at 1.2 [59].

When compared to WT vehicle-treated mice, both 3xTgAD (AD) and 3xTgAD/Polβ (ADP) models showed upregulation of clusters A-D and downregulation of cluster C, with our ADP model repeatedly showing greater deviation from WT mice (Fig. 5h). Our mice data are consistent with other AD brain transcriptomes and various mouse models, wherein cluster B genes are upregulated, and cluster C genes are predominately downregulated [59]. The response to UA was consistent in both AD and ADP mice as UA induced downregulation of clusters A-D and upregulation of cluster C. Notably, many cluster B and D terms were normalized by UA treatment in AD mice, whereas they were just mildly downregulated in ADP mice. Aging is the strongest risk factor for AD and cluster B terms are associated with immune function and are activated by age and in several other neurodegenerative and neuropsychiatric disorders [59]. In contrast, oligodendroglial-enriched cluster D terms, FPblue and TCXyellow, which are activated in select AD models, but rarely in other neurodegenerative disease models [59], were consistently downregulated by UA, suggesting that UA may be targeting multiple mechanisms of pathology and modulate both general immune, neuronal processes, and AD-specific pathophysiological pathways.

With respect to the AD neurodegenerative pathways and processes, most of the significant terms were upregulated in both AD and ADP models relative to WT mice, and again our ADP model showed greater deviation from WT than AD (Fig. 5i). UA treatment induced downregulation of myelination, cytokines, angiogenesis, tissue integrity, and growth factor signaling in both genotypes. Vesicle trafficking, trophic factors, and transcription and splicing were downregulated terms in the AD models relative to WT mice. UA caused upregulation of trophic factors and vesicle trafficking terms. Interestingly, UA downregulated oxidative stress in AD mice, but there was no change in ADP mice. Based on gene expression findings and behavioral studies [29, 30, 39], ADP mice possess more advanced AD pathology, and perhaps UA is less efficacious in advanced disease states.

Since there are only 760 genes on the NanoString AD panel, we collected all the genes from each pairwise comparison with a p-value of ≤ 0.05 and compared them in a three-way Venn diagram (Fig. 5j). There were 155 genes within the AD strain and 245 genes within the ADP strain comparisons. A majority of the genes in each pairwise comparison was unique to that comparison. To identify genes with a common UA response in AD and ADP, we combined the gene lists from the two strains and then selected those genes that were significantly changed (a fold change ≥ |1.2| in any comparison). This limited the gene list to 46, which is insufficient to do meaningful pathway analysis. Thus, we then sought to identified genes that were similarly significantly changed in the AD_WT or ADP_WT comparisons that were normalized by UA treatment (Fig. 5k). UA downregulated plasma membrane calcium transporting ATPase 1, *Atp2b1;* AF4/FMR2 family member 1, *Aff1*; ras homolog gene family member U, *Rhou*; dysbindin domain containing 2, *Dbndd2*; and zinc finger E-box binding homeobox 2, *Zeb2.* UA also upregulated carbonic anhydrase 10, *Car10*; nerve growth factor inducible, *Vgf*; ligand of numb-protein X 1*, Lnx1*; and cAMP-dependent protein kinase type I regulatory subunit Beta, *Prkar1b*. Among these nine genes, dysregulation of calcium in neurodegenerative disorders is common and *Atp2b1* removes calcium ions from eukaryotic cells. Further, ATP2B1 was previously identified as a hub gene in a transcriptomics study of human AD postmortem prefrontal cortical tissue [60]. VGF is also notable because a recent study identified VGF as a “high-confidence master regulator of AD associated networks” [61]. It is downregulated in human AD patients [62] and its expression correlates with disease progression. Similarly, it was down regulated in our mice and notably upregulated by UA. Overexpression in the 5xFAD mouse model partially rescued AD pathologies including memory impairment [61]. How this short list of genes contributes to the altered AD pathology is beyond the scope of the study here but warrants further study.

### 3.8 UA decreases neuroinflammation, mitochondrial markers, and DNA damage

UA is the most active, effective gut metabolite shown to stimulate mitophagy and acts as a potent anti-inflammatory and antioxidant agent [63]. To further investigate the beneficial effects of UA, the expression levels of important markers of mitochondrial stress, neuroinflammation, and DNA damage were explored in the hippocampus from Veh- and UA-treated 3xTgAD (AD) and 3xTgAD/Polβ^+/-^ (ADP) mice (Fig. 6a-g). In the absence of stress, PINK1 is constitutively imported, and undergoes cleavage by the mitochondrial processing peptidase and presenilin-associated rhomboid-like protease in the inner mitochondrial membrane [64, 65]. Upon mitochondrial damage, PINK1 is no longer imported; full-length PINK1 accumulates on the outer mitochondrial membrane and leads to autophosphorylation, which promotes kinase activation and facilitates binding to substrates Parkin and ubiquitin [66, 67]. As shown in Fig. 6a, our results showed that the accumulation of full-length Pink1 and Parkin were markedly increased compared with the WT group, accompanied by a significant increase in Bax protein expression in the hippocampus of AD and ADP mice. Their protein expressions were notably normalized by UA treatment, except for the Parkin level in AD mice (Fig. 6b, c, and d). We next examined neuroinflammation-related protein expression to evaluate the effects of UA on neuroinflammation. NLRP3 and Absent in Melanoma 2 (AIM2) inflammasome complexes are critical components of the innate immune system that mediate caspase-1 activation and induce pro-inflammatory cytokine production [68]. After activation of the NLRP3 inflammasome, the secretion of inflammatory cytokines such as IL-1β and TNFα leads to a strong inflammatory response, impairs nerve cell function, and consequently leads to cognitive dysfunction [68]. AIM2 is also a critical inflammasome sensor that recognizes cytosolic double-stranded DNA (dsDNA) via its HIN200 domain [69] and is responsive to microglial DNA damage [70]. Recent studies report that activation of the NLRP3 and AIM2 inflammasome signaling pathways play an important role in the neuroinflammation that drives AD pathology [70, 71]. Specifically, the deletion of the AIM2 inflammasome in AD mice models promoted dendrite branching, synaptic plasticity, and improvement in spatial memory [72]. Expressions of NLRP3 and AIM2 were significantly higher in AD mice than in WT and decreased strikingly after UA treatment in the hippocampus of AD and ADP mice (Fig. 6e and f). We also evaluated the phosphorylation of STAT3 on tyrosine 705 in both AD and ADP mice. STAT3 phosphorylation is critical for cytokine secretion and is linked to neuroinflammation in AD, and to IL-6 and TNFα [73]. Phosphorylation of STAT3 on Tyrosine 705 in the hippocampus was greatly elevated in both APP/PS1 transgenic mice [74] and in AD patients [75], and also verified in our APP/PS1 mouse brains (supplementary Fig 4d and 4f). As expected, the ratio of phosphorylated to total STAT3 protein was elevated in the hippocampus of AD and ADP mice compared with WT mice and was significantly reduced after UA treatment (Fig. 6g), suggesting that UA has beneficial effects on neuroinflammation. Several studies have provided strong evidence of crosstalk between DNA damage and inflammation [76]. Thus, we investigated the expression of γH2AX, a molecular marker of DNA damage and repair, to evaluate the effects of UA on DNA damage. As previously reported [11, 30], we showed that in the hippocampus of ADP mice, γH2AX was significantly increased compared with the WT group. Notably, after UA treatment, the AD mice showed a decrease in the protein level of γH2AX (Fig. 6h and i), as well as decreased PARylation and p-Parkin (Fig. 6h and supplementary Fig. 5g-h), indicating decreased DNA damage and/or increased DNA repair.

**Figure 6.**
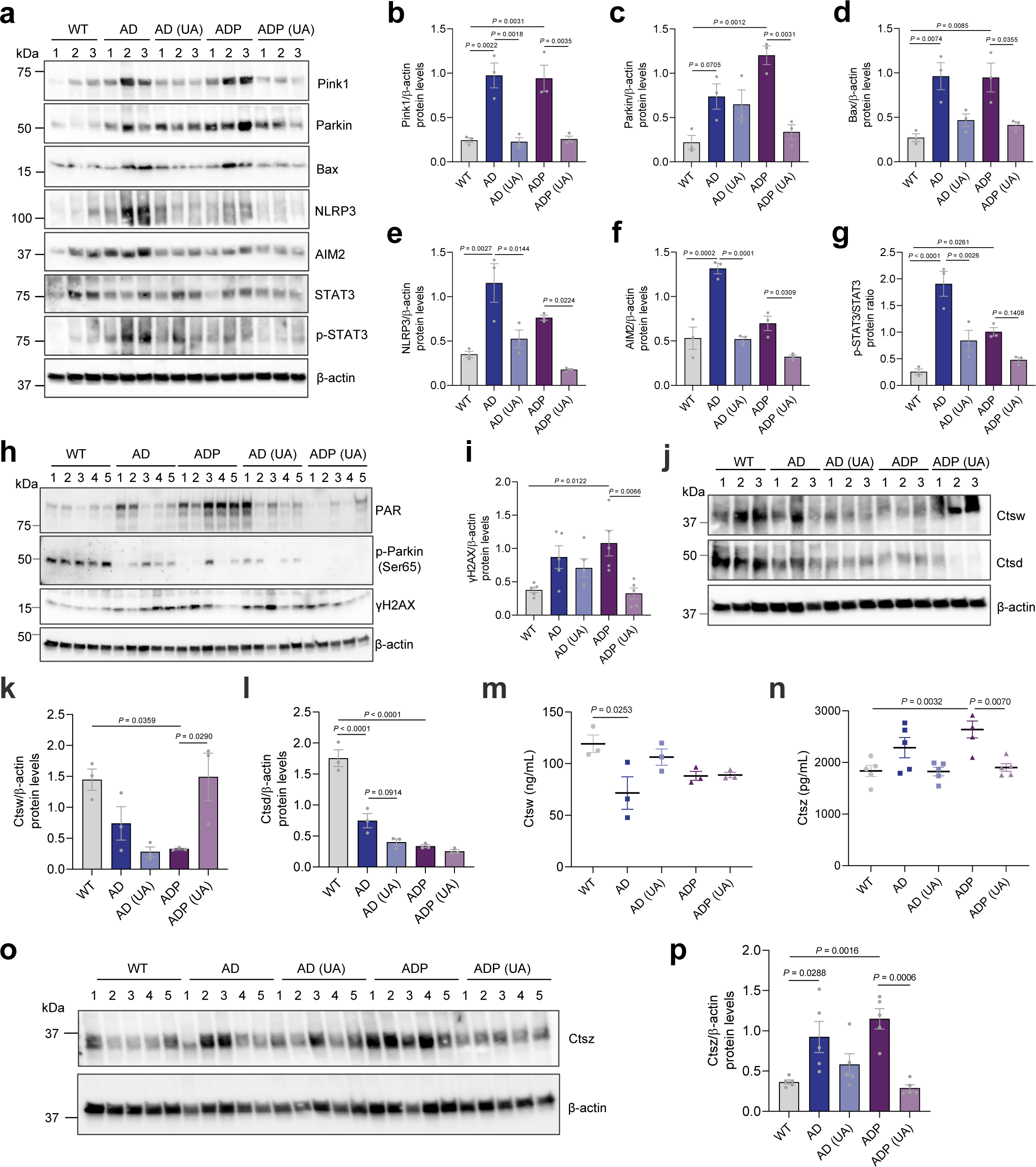
Effect of UA on mitochondrial dysfunction, neuroinflammation, and DNA damage. (**a**) Representative western blots showing PINK1, Parkin, STAT3, p-STAT3, NLRP3, AIM2, Bax in the hippocampus of WT, AD, and ADP mice with/without UA, n = 3 mice per group (1 F and 2 M). (**b–g**) Quantitation of protein levels for indicated proteins in Fig 6A. (**h-i**) Representative western blots showing PAR, p-Parkin, and γH2AX expression in the hippocampus and γH2AX quantification, n = 5 mice per group. (**j-l**) Representative western blots showing Ctsw and Ctsd and the quantitation of these protein levels in the hippocampus of WT, AD, and ADP mice with/without UA, n = 3 mice per group (1 F and 2 M). (**m-n**) Ctsz and Ctsw levels by ELISA assay. (**o-p**) Representative western blots showing Ctsz and the quantitation of the protein levels in the hippocampus of WT, AD, and ADP mice with/without UA, n = 5 mice per group. For h and o, WT (2 M & 3 F), AD (3 M & 2 F), AD+UA (3 M & 2 F), ADP (3 M & 2 F), and ADP+UA (2 M & 3 F). F, female, M, male. All bands were normalized to their respective loading controls β-actin. Error bars represent the mean ± SEM. Data were analyzed by one-way ANOVA analysis of variance with Tukey’s multiple comparisons test (**b-g**, **I**, **k-n**, **p**).

**Figure 7.**
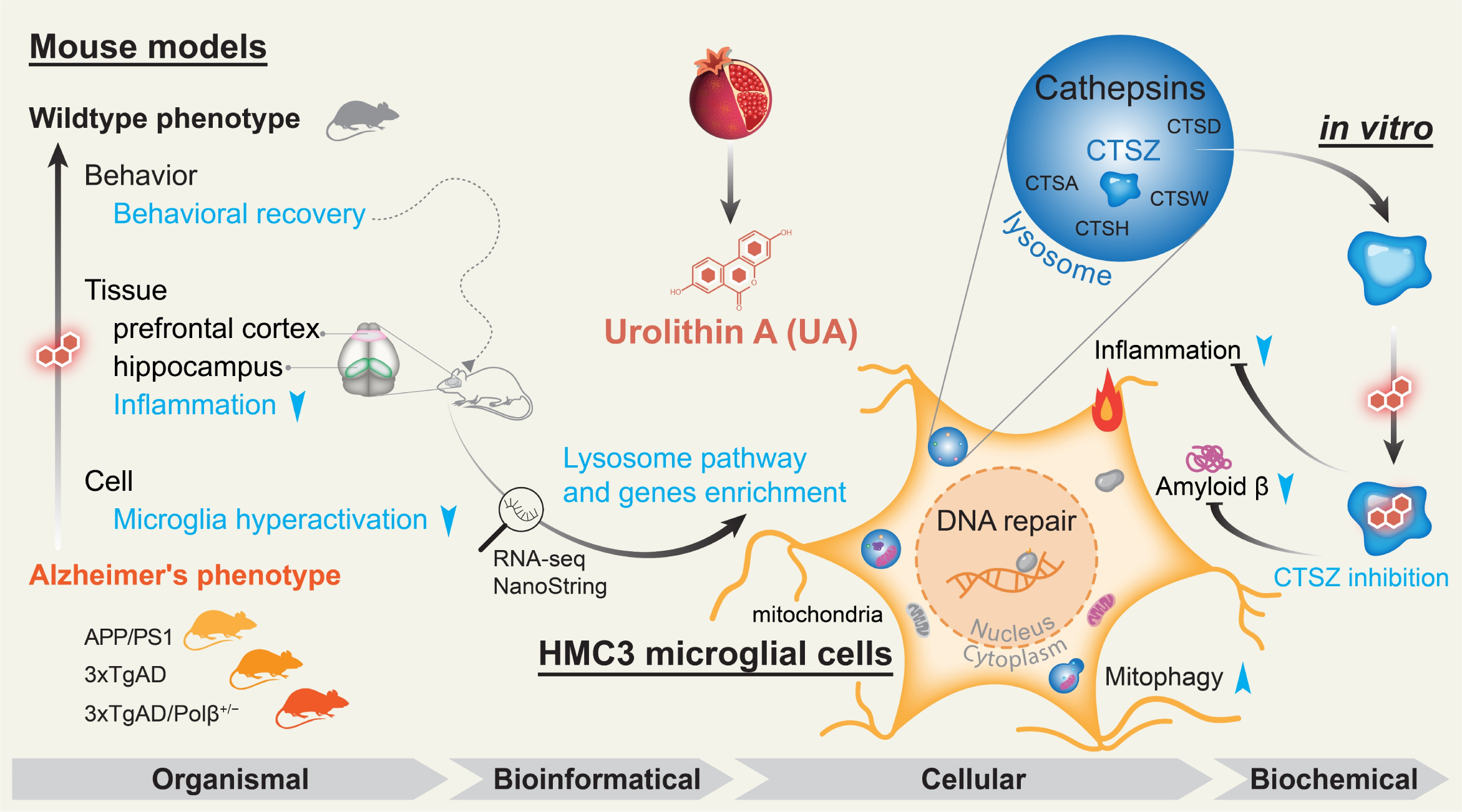
Graphical abstract. From organismal, bioinformatical, cellular and biochemical aspects, the effects and mechanisms of Urolithin A (UA) were investigated. We used different mouse models (including APP/PS1, 3xTgAD and 3xTgAD/Polβ^+/-^), and found that UA treatment improve behaviors, decreased inflammation and microglia hyperactivation. By using RNA-seq and NanoString, lysosomal pathways and gene were enrichment after UA treatment. RNA and protein levels of some lysosomal cathepsins are altered by UA treatment. We propose that UA inhibits CTSZ activity, to decrease Aβ and inflammation and induces mitophagy.

To further understand the pathway and gene changes related to mitochondria and metabolism in AD and ADP mice after UA, we used the NanoString metabolism panel, and found some significantly changed terms, including lysosomal degradation and NF-κB terms, as well as several cathepsins and the *Ctsz* gene (supplementary Fig. 6e-f). We also validated the expression of lysosomal Ctsw, Ctsd, and Ctsz in the hippocampus of AD and ADP mice models, respectively. The Ctsw and Ctsd were lower in AD mice than in WT, and UA treatment did not notably affect their expression, except for the Ctsw levels in ADP mice (Fig. 6j-m). The level of Ctsz expression was significantly elevated in AD brains and normalized after UA treatment (Fig. 6n-p). Similar results were seen in APP/PS1 mice, so across multiple mouse models, we find cathepsins and in particular Ctsz, regulated by UA.

## 4 DISCUSSION

Our study identifies mechanisms underlying the beneficial effects of long-term UA treatment in AD transgenic mice models. APP/PS1 mice treated long-term with UA took significantly less time to reach the platform, similar to WT (Fig. 1b), We find that UA treatment has beneficial effects on multiple aspects of neuropathology (long-term potentiation, Aβ, pTau, neuroinflammation, lysosomal function and DNA damage) in the hippocampus and PFC regions, which play central roles in AD. As shown in the above figures, key drivers of the neuroinflammatory process including IL-1β, NLRP3, AIM2, and STAT3 were higher in the brains of AD mice than in WT and decreased notably after UA treatment. Importantly, we are the first to present evidence that Ctsz plays an important role in UA induced therapeutic effects for AD. Thus, our results are consistent with literature reporting that UA has therapeutic properties in AD mice brains [11, 21]. We have summarized a table comparing UA treatment for 1∼2 months (which we reported previously [11]) and UA treatment for 5 months in AD mouse models (Supplementary Table 1). We have now compared and summarized the behavioral improvement (including cognitive function, smelling, etc.), the improvement of pathological characteristics of AD (including Aβ, tau, etc.), the neuroinflammation, the improvement of mitophagy, and other aspects of each AD mouse treated with 1∼2-month UA and 5-month UA. Through this detailed comparison, we provide information for future translational research.

UA is thought to play a critical role in supporting mitochondrial function in neuronal dysfunction and neurodegeneration [11, 21]. Mitochondrial dysfunction has been widely reported in studies of AD patients and AD models [5]. Notably, we found that full-length PINK1 and Parkin recruitment were significantly increased in AD brains with mitochondrial damage likely leading to activated Bax (Fig. 6). Damage to mitochondria results in activation of mitophagy and affects the mitochondrial outer membrane permeabilization (MOMP) that in turn triggers apoptotic cell death. Upon the collapse of the mitochondrial membrane potential (Δψm), the import and degradation of PINK1 are blocked, and PINK1 accumulates on the outer mitochondrial membrane [66, 67]. Also, Bax translocates from the cytosol to the mitochondrial membrane, thereby facilitating the release of pro-apoptotic proteins during apoptosis. UA treatment of AD and ADP mice normalized the levels of these proteins to WT.

UA has also been shown to have a positive effect on DNA damage and repair in AD [11]. DNA damage and DNA repair deficiency can lead to neuronal dysfunction in aged brains and in the brains of AD patients and transgenic mice [30, 77]. In our study, DNA damage was identified by high levels of γH2AX in the hippocampus of brain tissue and these decreased after treatment with UA (Fig. 6h). A previous study also reported that UA suppressed DNA double strand breaks (DSBs) in bone marrow-derived macrophages [78]. These finding suggest that treatment with UA may regulate DNA damage through decreasing the amount of oxidative DNA damage and mitochondrial oxidative stress or by increasing DNA repair.

UA at dose from 250 mg to 2000 mg in humans [26] and 1-450 mg/kg in mice [79] has been reported to be safe. UA increased muscle strength and physical performance in a 6 min walk test in elderly humans after four months of supplementation [80]. Other studies reported that UA improved motor activity in the rotarod test and increased total distance traveled and average speed in the open field test in young C57BL/6J mice [81] and in 3xTg AD mice [82]. However, we noted that UA decreased the distance travelled by 3xTgAD mice in the open field test (Fig. 1l), while the open field activity of APP/PS1 mice in our study were normal compared with age-matched control mice (supplemental Fig. 1a-d). The mice were of different ages in these two cohorts. The APP/PS1 mice were 7 months old and the AD and ADP mice were 17 months old at testing. Another potential factor is that 3xTgAD mice express mutant tau whereas the APP/PS1 mice do not express the transgene. Additionally, another difference between the strains was sex, the APP/PS1 mice were all male while the 3xTgAD used are both sexes. Combined there are a number of potential reasons why the strains have a different response.

Growing evidence implicates the important roles for lysosomal function in AD development Particularly, alterations in lysosomal cathepsins in the central nervous system (CNS) contribute to the pathogenesis of neurodegenerative diseases as seen for AD, as well as synucleinopathies (PD and Dementia with Lewy Body), and HD [83]. In our study, one of the most significant findings was that Ctsz was highly expressed in multiple AD transgenic mouse models and its expression was normalized by UA treatment (Figs. 4 and 6), suggesting a critical role of Ctsz in UA-induced therapeutic effects for AD. Further analysis also revealed the mutual functions between UA and CTSZ. Inhibiting Ctsz activity reduced the Aβ_42_ level in the MHC3 cells and also compromised UA increased lysosomal functions, maybe because of Ctsz inhibition decreased Aβ_42_ levels, suggesting that UA is partly dependent upon Ctsz to facilitate lysosomal activities. Over the past decade, increasing evidence has emerged implicating Ctsz (also known as cathepsin X/P) in the inflammatory processes leading to neurodegeneration. Ctsz has been shown to be highly expressed and is secreted by microglia and astrocytes in response to neuronal damage and inflammatory stimulus, both *in vitro* and *in vivo* [84, 85]. Unlike other cathepsins, such as Ctsa, Ctsh, and Ctsw, Ctsz is unique in its ability to modulate NLRP3-inflammasome activation [86]. It is widely expressed early in CNS development and linked to IL-18 and IL-1β production [49, 50]. In particular, high levels of CTSZ have been observed in degenerating brain regions of the ALS transgenic mouse model and in the post-mortem cortex tissues from AD individuals [87, 88]. In the transgenic APP/PS1 mouse model, Ctsz upregulation has been observed in microglial cells surrounding amyloid plaques, suggesting that activated microglia may have contributed to inflammation-induced neurodegeneration by secreting Ctsz [87]. In addition, a decreased neuroinflammatory state was observed in a *Ctsz* knockout mice with experimental autoimmune encephalomyelitis [49], supporting that Ctsz may be a potentially interesting therapeutic target in AD neuroinflammation.

UA-mediated inhibition of neuroinflammation and lysosomal dysfunction may contribute to the improvement of AD pathophysiology. Our results provide the first experimental evidence that long-term UA treatment may ameliorate neuroinflammation by regulating Ctsz expression in AD transgenic mouse models.

## Abbreviations

AD: Alzheimer’s disease
UA: Urolithin A
Ctsz: Cathepsin Z
Aβ: Amyloid β
pTau: phosphorylated Tau
NR: nicotinamide riboside
PINK1: PTEN-induced kinase 1
PDR1: pleiotropic drug resistance 1
DCT1: DAF-16/FOXO-controlled germline-tumor affecting-1
ADP: 3xTgAD/Polβ+/-
PFC: prefrontal cortex
GFAP: Glial fibrillary acidic protein
SIRT3: sirtuin 3
Ctsa: Cathepsin A
Ctsh: Cathepsin H
Ctsl: Cathepsin L
Ctss: Cathepsin S
IL-1β: Interleukin-1β
NLRP3: NLR Family Pyrin Domain Containing 3
Lamp1: lysosomal-associated membrane protein 1
DqBSA: Dye Quenched-Bovine Serum Albumin
Baf: bafilomycin
HD: Huntington’s disease
MCP-1: Monocyte chemoattractant protein-1
MIP-1α: Macrophage Inflammatory Protein-1 Alpha
TNFα: tumor necrosis factor
KC: keratinocyte-derived cytokine
TREM2: triggering receptor expressed on myeloid cell 2
TYROBP: tyrosine motif binding protein
AIM2: Absent in Melanoma 2

## supplementary Figure legends

**sFigure 1.**
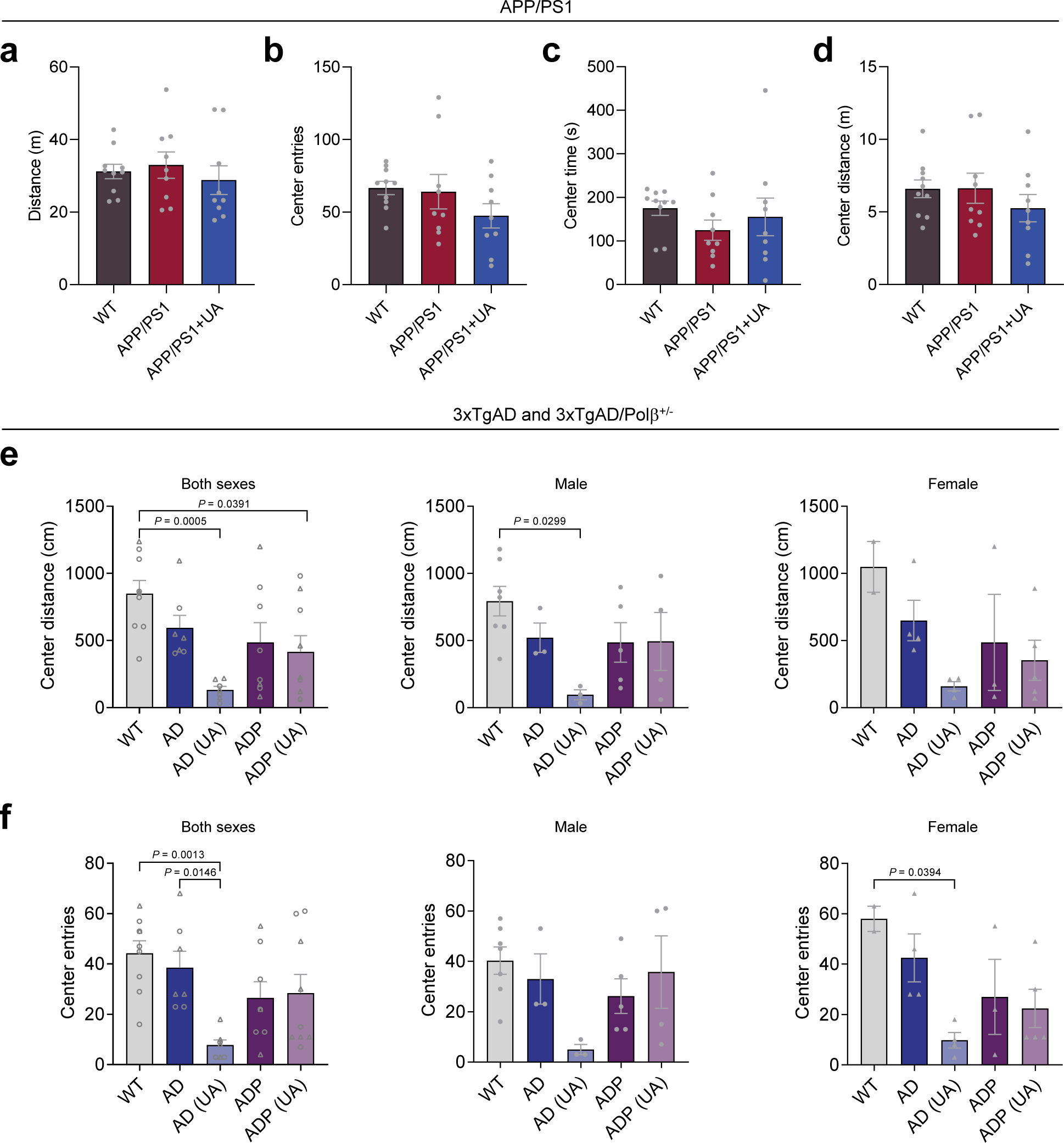
UA didn’t change motor function in AD mice. (**a-d**) The open field test of WT, APP/PS1 and UA-treated APP/PS1 mice, n = 9-10 male mice. (**e-f**) The open field test of WT, AD, ADP and UA-treated AD and ADP mice, n = 7-9 mice. The right two graphs are analysis by different sexes. Data were analyzed by one-way ANOVA with Tukey’s multiple comparisons test (**a-f**). Data were shown as mean ± SEM. For a-d, all mice used are males. For e-f, WT (7 M and 2 F), AD (3 M and 4 F), AD + UA (3 M and 4 F), ADP (5 M and 3 F), ADP + UA (4 M and 5 F). F, female, M, male.

**sFigure 2.**
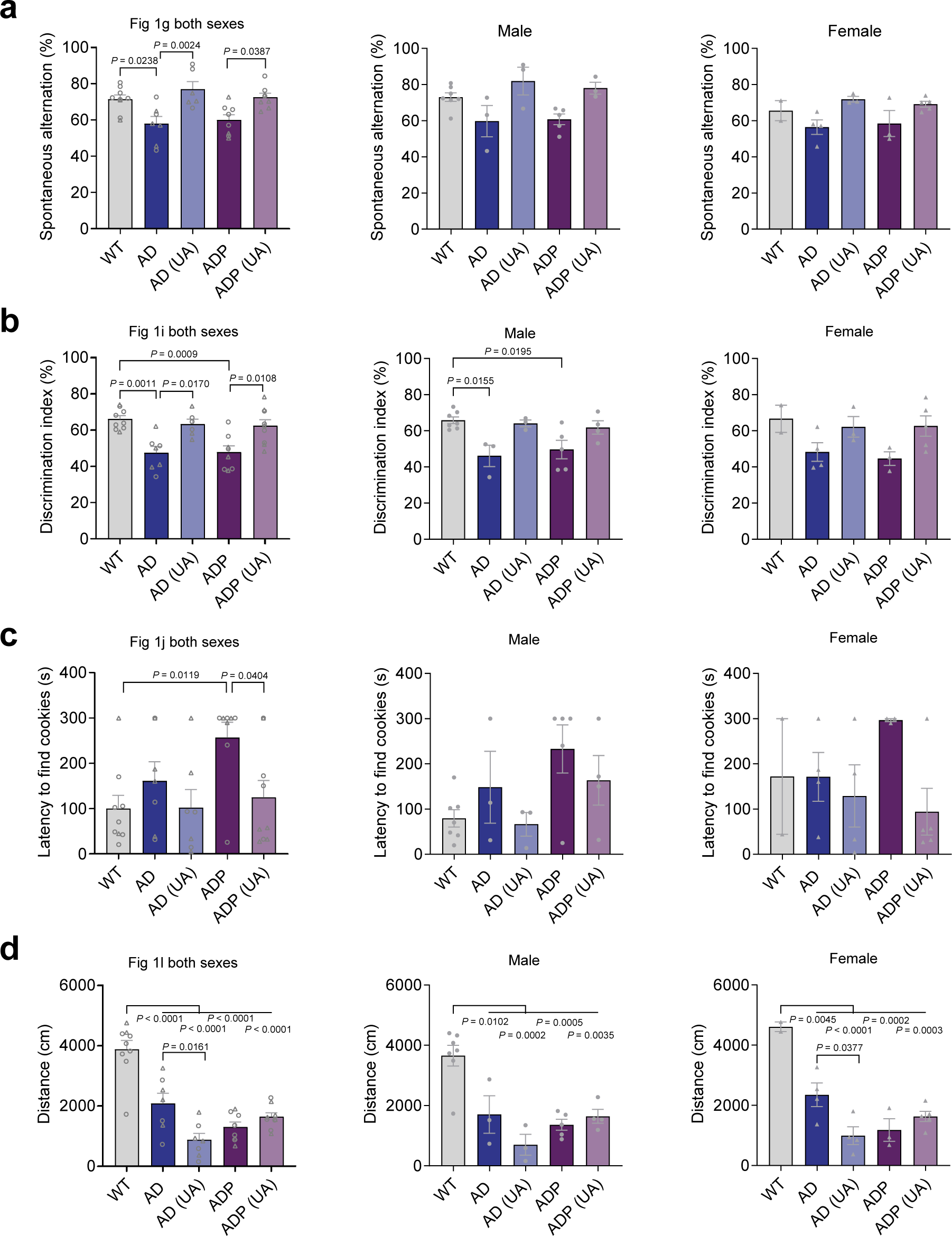
Behavioral analysis by different sexes. (**a**) The Fig. 1g Y-maze analysis by male and female sexes. (**b**) The Fig. 1i object recognition test analysis by male and female sexes. (**c**) The Fig. 1j smelling test analysis by male and female sexes. (**d**) The Fig. 1l open field test analysis by male and female sexes. Data were analyzed by one-way ANOVA with Tukey’s multiple comparisons test (**a-d**). Data were shown as mean ± SEM. For a-b, WT (7 M and 2 F), AD (3 M and 4 F), AD + UA (3 M and 3 F), ADP (5 M and 3 F), ADP + UA (3 M and 5 F); for c-d, WT (7 M and 2 F), AD (3 M and 4 F), AD + UA (3 M and 4 F), ADP (5 M and 3 F), ADP + UA (4 M and 5 F). F, female, M, male.

**sFigure 3.**
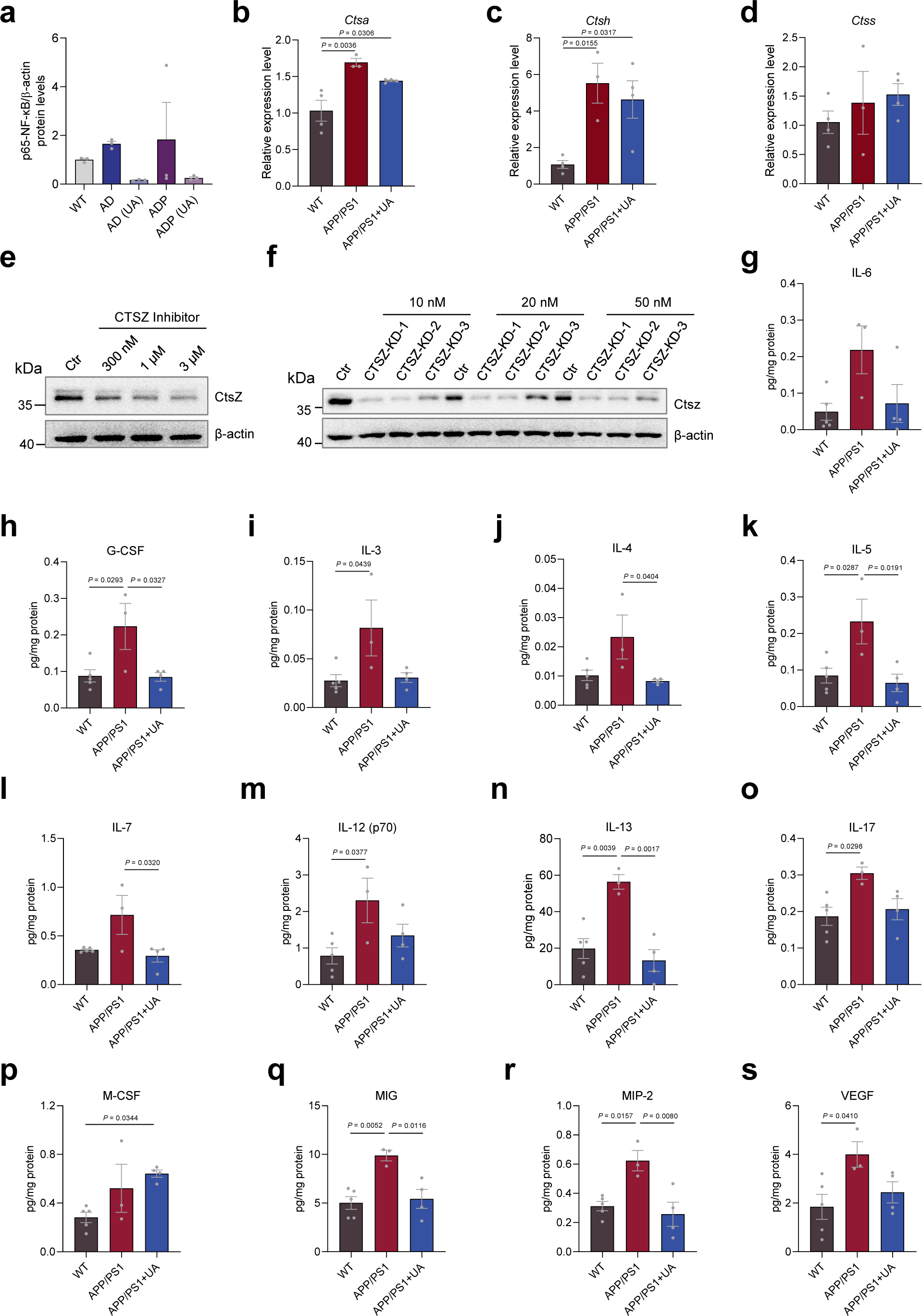
UA decreased neuroinflammation in AD mice brains. (**a**) Quantification of protein levels for p65-NF-κB. n = 3 mice per group. All samples were normalized to respective loading controls β-actin. (**b-d**) qPCR analysis for the relative expression of *Ctsa* (a), *Ctsh* (b) and *Ctss* (c), n = 3-4 mice. (**e-f**) The CTSZ inhibitor and two si-*CTSZ* efficiency detected by western blot in HMC3 cells. (**g-s**) The cytokine assay of cytokine levels in in WT, APP/PS1 and APP/PS1 + UA mice cortex lysates, including IL-6, G-CSF, IL-3, IL-4, IL-5, IL-7, IL-12 (p70), IL-13, IL-17, M-CSF, MIG, MIP-2, VEGF, n = 5 in the WT group; n = 3 in the APP/PS1 group; n = 4 in the APP/PS1 + UA group. Data were shown as mean ± SEM. Data were analyzed by one-way ANOVA with Tukey’s multiple comparisons test (**a-d**, **g-s**). For APP/PS1 strain mice, all used are males. For AD, AD+UA: 2 males and 1 female; For WT, ADP and ADP+UA: 1 male and 2 females.

**sFigure 4.**
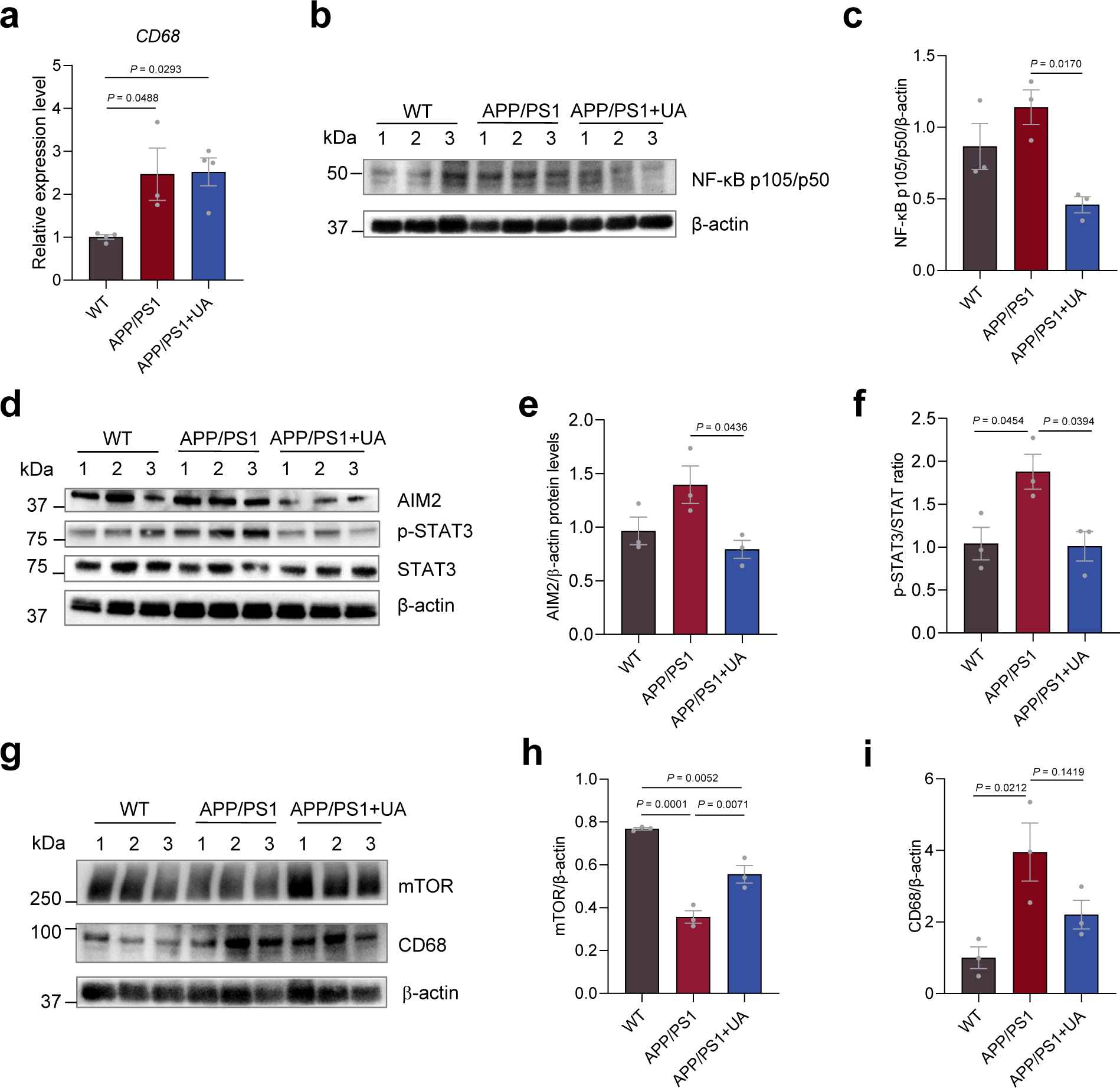
UA decreased neuroinflammation in the hippocampus of AD mice brain lysates. (**a**) qPCR analysis for the relative expression of *CD68*, n = 3-4 male mice. (**b**) Western blots of NF-κB p105/p50, n = 3 male mice per group and (**c**) quantitation in (b). (**d**) Western blots of the indicated proteins, n = 3 male mice per group. (**e-f**) Quantitation of (**d**). (**g**) Western blots of the indicated proteins, n = 3 male mice per group. (**h-i**) The quantitation of mTOR and CD68 protein levels in (g). Data were analyzed by one-way ANOVA with Tukey’s multiple comparisons test (**a**, **c**, **e-f, h-i**). Data were shown as mean ± SEM. For APP/PS1 strain mice, all used are males.

**sFigure 5.**
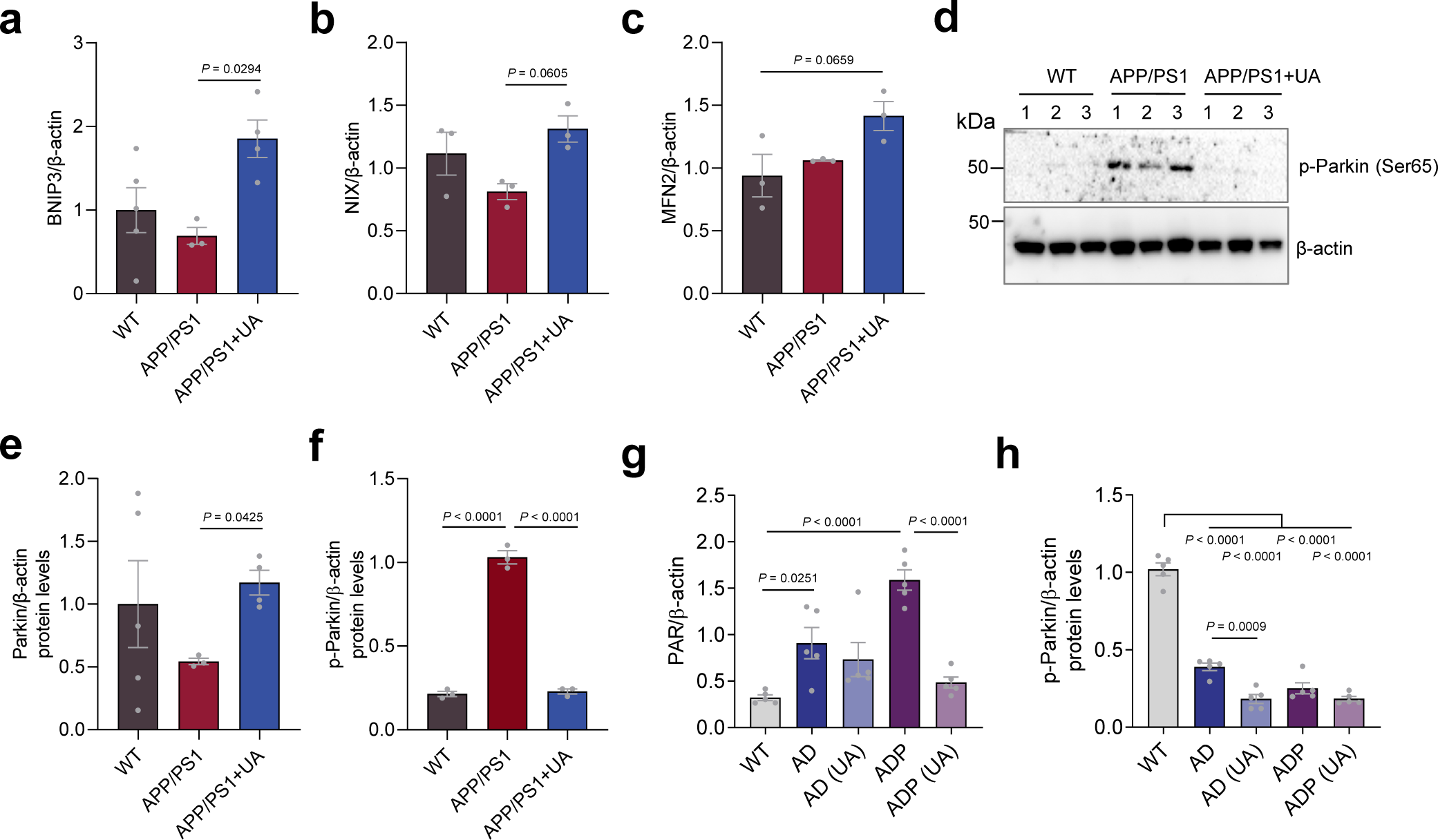
UA induced mitophagy, increased mTOR level and DNA damage. (**a-c**) Quantitation of the protein levels in Fig. 5d, n = 3 male mice. (**d-f**) Representative western blots showing p-Parkin and quantitation of Parkin (Fig. 5d) and p-Parkin protein levels in the cortex of WT, APP/PS1, and APP/PS1 with UA mice, n = 3 mice per group. (**g-h**) Quantitation of PAR, p-Parkin in the hippocampus of WT, AD, and ADP mice with/without UA in Fig. 6h, n = 5 mice per group. WT (2 M & 3 F), AD (3 M & 2 F), AD+UA (3 M & 2 F), ADP (3 M & 2 F), and ADP+UA (2 M & 3 F). F, female, M, male. Data were analyzed by one-way ANOVA with Tukey’s multiple comparisons test (**a-c**, **e-h**). Data were shown as mean ± SEM. For APP/PS1 strain mice, all used are males.

**sFigure 6.**
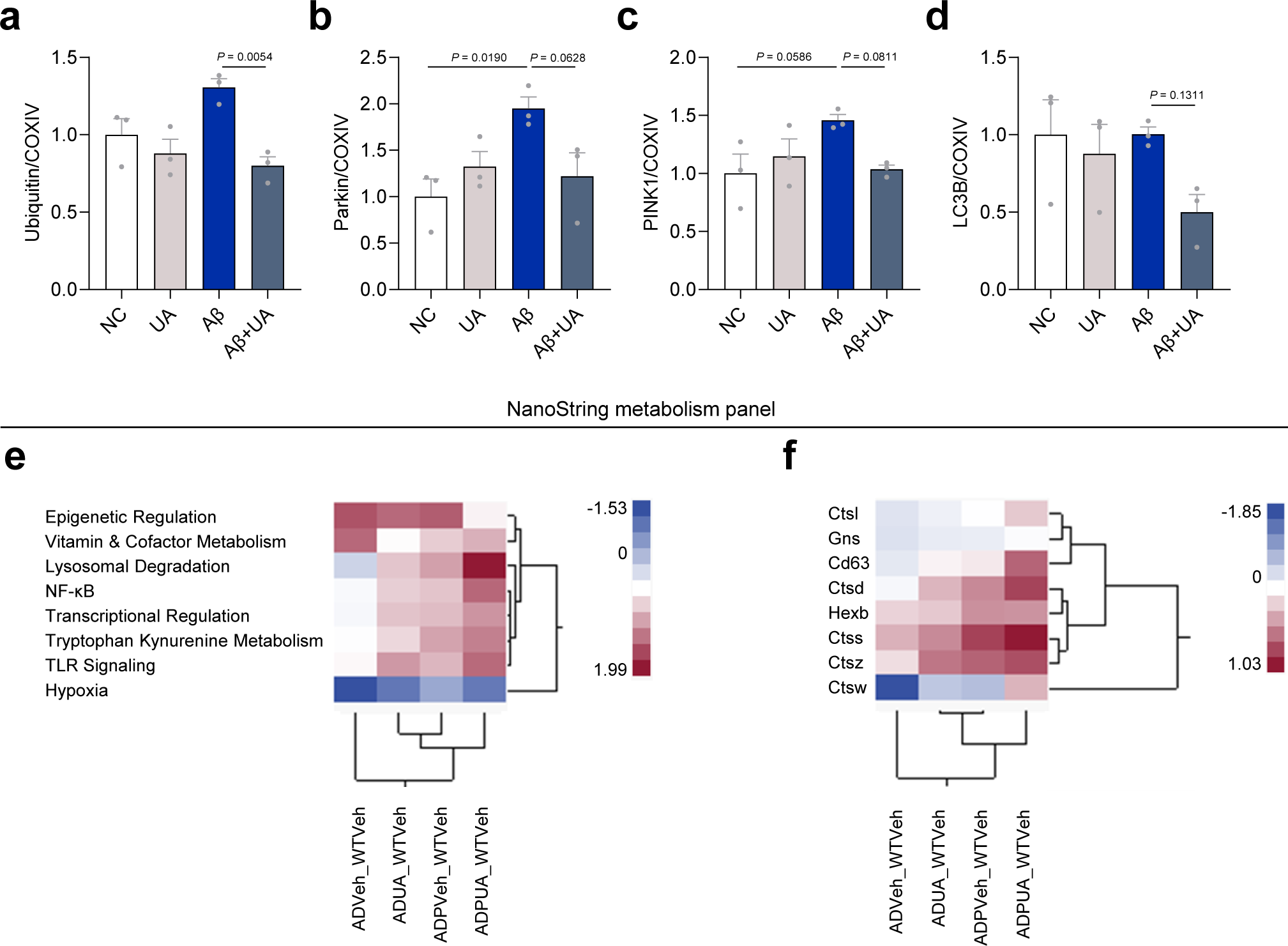
UA normalized Aβ-induced mitophagy-related proteins in mitochondrial extracts, and the NanoString analyses in AD and ADP mouse brains. (**a-d**) The quantification of Fig. 5g. n = 3 independent repeats. (**e**) NanoString metabolism panel analysis reveals lysosomal term changes. n = 4 for WT and n = 5 mice per group for the rest. WT (2 M & 2 F), AD, AD+UA, and ADP (3 M & 2 F), ADP+UA (2 M & 3 F). F, female, M, male. (**f**) Subset of significantly changed lysosomal genes identified in NanoString metabolism panel analysis.

**sTable 1.**
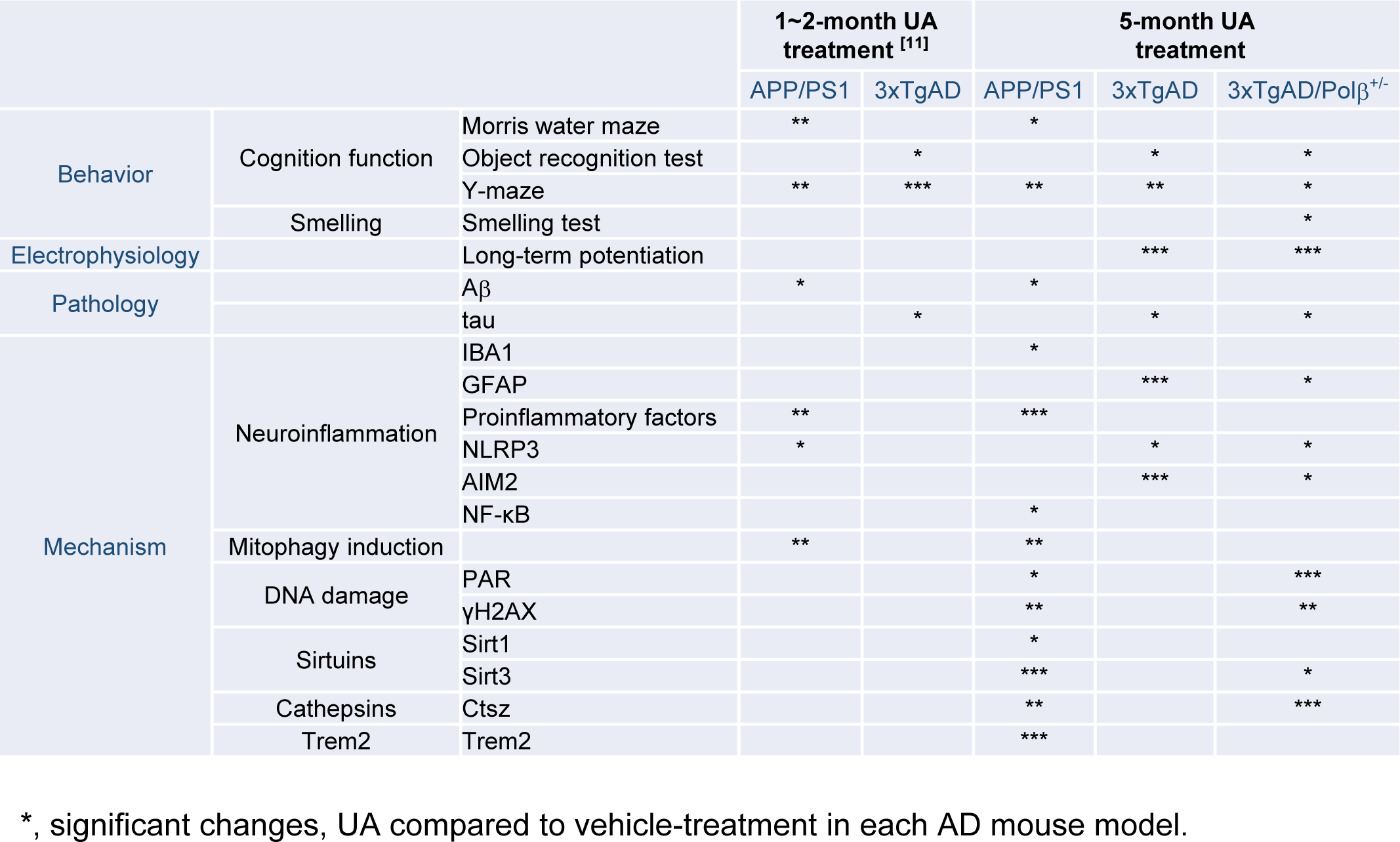
Comparison between 1∼2-month UA treatment and 5-month UA treatment in AD mouse models.

## Acknowledgements

We thank Y.Q. Zhang, E. Lehrmann, and K. Becker for the array data; J. Tian for the technical support with the experiments.

## Authors’ contributions

Y.H. and V.A.B. designed the experiments. Y.H., B.Y., and Y.W. performed the animal treatment and behavior tests. Y.H. performed the microarray and D.L.C. analyzed the microarray data. X.C. and J.P. performed the western blot, ELISA, and NanoString experiments, and D.LC. and J.P. analyzed NanoString results. Q.Z., M.H., H.B.M. performed the lysosomal assay and immunofluorescence assays. Z.L. performed the *in vitro* assay. Y.W. performed the electrophysiology experiments. E.F.F. contributed to constructive discussions and editing of the manuscript. Y.H., X.C., J.P., D.L.C., and V.A.B. wrote the manuscript.

## Declarations

### Ethics approval and consent to participate

All animal experiments were performed and approved by the National Institute on Aging (NIA) Animal Care and Use Committee, and met all relevant ethical regulations (study protocol number 361-ODS-2023).

### Consent for publication

Not applicable.

